# Assortative mating, sexual selection and their consequences for gene flow in *Littorina*

**DOI:** 10.1101/2020.01.28.922658

**Authors:** Samuel Perini, Marina Rafajlović, Anja M. Westram, Kerstin Johannesson, Roger K. Butlin

**Author notes:** Correspondence: Samuel Perini < >. These authors contributed equally to this work (Samuel Perini, Marina Rafajlović).

## Abstract

When divergent populations are connected by gene flow, the establishment of complete reproductive isolation usually requires the joint action of multiple barrier effects. One example where multiple barrier effects are coupled consists of a single trait that is under divergent natural selection and also mediates assortative mating. Such multiple-effect traits can strongly reduce gene flow. However, there are few cases where patterns of assortative mating have been described quantitatively and their impact on gene flow has been determined. Two ecotypes of the coastal marine snail, *Littorina saxatilis*, occur in North Atlantic rocky-shore habitats dominated by either crab predation or wave action. There is evidence for divergent natural selection acting on size, and size-assortative mating has previously been documented. Here, we analyze the mating pattern in *L. saxatilis* with respect to size in intensively-sampled transects across boundaries between the habitats. We show that the mating pattern is mostly conserved between ecotypes and that it generates both assortment and directional sexual selection for small male size. Using simulations, we show that the mating pattern can contribute to reproductive isolation between ecotypes but the barrier to gene flow is likely strengthened more by sexual selection than by assortment.

---

The formation of new species requires the evolution of reproductive isolation through the accumulation of barriers to gene flow. Where divergence occurs in allopatry, different barrier effects are automatically associated, but with gene flow these associations need to be created and maintained by selection operating against the effects of recombination (Felsenstein 1981; Smadja and Butlin 2011). One example is the increase in the overall barrier to gene flow resulting from associations between divergent selection and assortative mating (Kirkpatrick and Ravigné 2002; Gavrilets 2004; Sachdeva and Barton 2017). If this requires the build-up of linkage disequilibrium among separate sets of loci controlling divergently selected traits, signal traits and preferences, it may be easily opposed by gene flow and recombination (Servedio 2009; Smadja and Butlin 2011). However, some types of traits and forms of assortative mating reduce the number of associations that need to be maintained and so are expected to be more likely to contribute to reproductive isolation. ‘Multiple-effect’ traits are traits that contribute to more than one barrier effect (Smadja and Butlin 2011). Sometimes, the term ‘magic trait’ is used for a subset of multiple-effect traits where the trait under divergent selection also contributes to assortative mating (Servedio et al. 2011).

Assortative mating might depend on a matching rule where there is no separation of signal and preference and so they necessarily coevolve. Alternatively, it might depend on a preference/trait rule where signal and preference are separate and coevolution is not guaranteed (Kopp et al. 2018). In the extreme, there might be only a single trait involved, such as habitat choice or flowering time (“matching rule by a grouping mechanism”; Kopp et al. 2018; Servedio and Kopp 2012). The ecological trait is then a multiple-effect trait and no other trait is needed to generate assortment. Body size in *Gasterosteus* sticklebacks (McKinnon and Rundle 2002) is an example of a multiple-effect trait where mating is based on phenotypic similarity of a trait under divergent natural selection. Wing color-pattern in *Heliconius* butterflies (Merrill et al. 2014, 2019) is a multiple-effect trait that is also under divergent natural selection but contributes instead to assortative mating primarily through the signal component of a signal-preference system. Assortment can also be driven by the preference component, as in the case of cichlids where color sensitivity influences both foraging and mate choice (Seehausen et al. 1999).

The evolution of assortative mating, and the barrier to gene flow that it generates, can also be impacted by sexual selection. Assortative mating can occur without variation in mating success among individuals. However, behavioral interactions between males and females that generate assortative mating will often also generate sexual selection. For example, males with intermediate trait values might find mates with common, intermediate preferences more easily than males with extreme values, generating stabilizing sexual selection (Servedio et al. 2011; Servedio and Hermisson 2019). If the trait is also under natural selection and the ecological optima differ between populations, the stabilizing sexual selection may oppose divergence, but it can also contribute to reproductive isolation once divergence is achieved. Sexual selection must be divergent in order to contribute to the ongoing evolution of reproductive isolation but differences in preference between populations may not be enough: if, for example, preferences are less divergent than the traits on which they are based, sexual selection can lead to decreased differentiation between populations after contact (Servedio and Boughman 2017). There are still few empirical studies that have demonstrated the extent to which sexual selection contributes to reproductive isolation or its ongoing evolution (Maan and Seehausen 2011; Servedio and Boughman 2017).

Whatever the nature of assortative mating and sexual selection, it is important to quantify their contribution to the overall barrier to gene flow during the process of speciation. The contributions of individual barriers can be estimated by breaking down reproductive isolation into its components (Coyne and Orr 2004 pp. 63–65; Lowry et al. 2008; Sobel and Chen 2014). In these calculations, the estimate of assortative mating typically comes from comparisons between divergent populations as indices of premating isolation [e.g., Yule’s V (Gilbert and Starmer 1985) and I_PSI_ (Rolán-Alvarez and Caballero 2000)]. In turn, these isolation indices come from experiments where individuals can mate either within their own population or with an individual from a divergent population (e.g., Matsubayashi and Katakura 2009). However, these indices risk over-simplifying the mating pattern and they fail to account for the presence of the intermediate phenotypes that occur whenever reproductive isolation is incomplete (Coyne and Orr 2004; Irwin 2019).

Hybrid zones provide excellent conditions for quantifying the extent to which gene flow between distinct populations is reduced by divergent natural selection and assortative mating (Hewitt 1988). In contact zones between divergent populations, hybrids can form and display a wide range of trait combinations (Barton and Hewitt 1985; Mallet 2005). For example, two locally adapted populations can evolve different trait values for a quantitative trait but a continuous cline in the trait will typically be maintained across the habitat boundary. Gene exchange will continue but will be impeded, particularly for loci contributing to selected traits and loci closely linked to them, with the width of the cline providing an estimate of barrier strength (Barton and Gale 1993). This provides an excellent opportunity to quantify the barrier effects of assortative mating and sexual selection. It has been argued that assortative mating based on clinally-varying traits will generate only a weak barrier to gene flow because individuals that meet one-another in the hybrid zone rarely differ much in trait values, allowing little opportunity for discrimination (Irwin 2019). This logic does not apply to traits with a very simple genetic basis because they are not expected to show a continuous cline across the habitat boundary. Selection resulting from the reduced fitness of hybrids can, in theory, increase reproductive isolation (reinforcement) but the conditions required are quite stringent (Liou and Price 1994; Price 2008; but see Servedio and Noor 2003). Both the barrier generated by assortative mating and the likelihood of reinforcement depend on the mechanism of assortment (Kopp et al. 2018) and the genetic architecture of the traits involved (Felsenstein 1981; Smadja and Butlin 2011).

To understand the impact of departures from random mating on the barrier to gene flow in a hybrid zone, it is necessary to quantify the mating pattern. By ‘mating pattern’, we mean the function that predicts the probability of mating, given an encounter between a male and female with specified phenotypes. This might vary across the zone. Given the mating pattern and the distributions of male and female phenotypes, it is possible to predict the strength of assortative mating and sexual selection at any point in the zone. In turn, this can be used to infer the barrier effect in a way that cannot be deduced from interactions between individuals from divergent, parental populations alone. The impacts of assortative mating and sexual selection can also be separated (Servedio and Boughman 2017).

Here, we address these issues in the marine snail *Littorina saxatilis*, combining extensive empirical data from mating experiments with a model-based quantitative description of the mating pattern that we then use to infer assortative mating and sexual selection in the field. We also use the mating pattern as an input to computer simulations to study the barrier effects of both assortative mating and sexual selection. *Littorina saxatilis* is an intertidal marine snail forming multiple ecotypes, facilitated by low dispersal due to direct development. The Wave and the Crab ecotypes (simply “Wave” and “Crab” in the following) are encountered widely in wave-exposed and crab-rich habitats, respectively, over the species’ North Eastern Atlantic distribution (Johannesson et al. 2010; Butlin et al. 2014). Wave individuals live on cliffs, and they have evolved a relatively large foot, thin shell, a bold behavior and small sizes, whereas Crab snails live among boulders, and differ from the Wave snails by a larger, thicker shell with a narrower foot, showing a wary behavior. Trait differences between ecotypes, such as size and shape of the shell, are the result of local adaptation, most likely induced by wave action in the wave-exposed habitat and crab predation in the crab-rich habitat (Johannesson 1986; Boulding et al. 2017; Le Pennec et al. 2017). Many genomic regions potentially involved in the divergence process in *L. saxatilis* have been identified, including several putative inversions (Westram et al. 2018; Faria et al. 2019; Morales et al. 2019).

Divergent natural selection is a powerful barrier against gene flow between Wave and Crab snail populations but there are also suggestions for other components of isolation such as habitat choice and size-assortative mating (Janson 1983; Rolán-Alvarez et al. 1997; Cruz et al. 2004; Johannesson et al. 2016). Assortative mating has been investigated in empirical studies both in the field and the laboratory showing that Crab and Wave ecotypes mate assortatively in sympatry (Yule’s V, I_PSI_ and *r_i_* values significantly different from random mating and as high as 0.96; Johannesson et al. 1995; Hull 1998; Rolán-Alvarez et al. 1999; Cruz et al. 2004; Conde-Padín et al. 2008) and that female and male sizes in field-collected mating pairs were highly correlated (Pearson correlation coefficients 0.3; Rolán-Alvarez et al. 1999, 2004; Johannesson et al. 1995). Assortment is accompanied by a component of sexual selection on size that favors large females and small males (Ng et al. 2019). Furthermore, copulation time, as well as distances that males follow female trails before mating, are longer for similarly sized pairs with the female being on average slightly larger than the male (Hollander et al. 2005; Johannesson et al. 2008). Because the average sizes of the ecotypes are very different (adult Crab snails are two to three times longer than adult Wave snails) this generates assortment among ecotypes, with little evidence for effects of traits other than size. Among littorinid snails of various species, males preferentially track and mate females slightly larger than themselves (‘similarity-like’ mechanism plus a constant; Erlandsson and Johannesson 1994; Saltin et al. 2013; Ng and Williams 2014; Fernández-Meirama et al. 2017; Ng et al. 2019) suggesting that this mating pattern is ancestral.

There is strong evidence for the presence of assortative mating by size in *L. saxatilis* plus the opportunity for sexual selection on size. Thus, size is a multiple-effect trait, under direct divergent selection between the Crab and Wave habitats and also a key trait influencing mating success. However, for the general reasons discussed above, it is unclear to what extent this assortative mating contributes to the barrier to gene flow between the two ecotypes where they meet in natural contact zones. It is also not known whether sexual selection enhances the reproductive barrier in this system. Hence, we asked what the barrier effect of size-assortative mating and sexual selection is in natural contact zones in these snails. First, we quantified the mating probability given encounters between snails with a wide range of sizes and shapes. Second, we used the resulting mating pattern to infer assortative mating and sexual selection across the contact zones between populations of the Crab and Wave ecotypes. Finally, based on these estimates of assortment and sexual selection, we assessed the likely barrier effects of these two components of isolation by performing individual-based computer simulations.

## Materials and Methods

### Sampling, Phenotypes and Mating Experiment

Along-shore transects including Crab-Wave contact zones were sampled on four small islands on the Swedish west coast. Each sampled transect was approximately 300 m long and included one boulder field (Crab snail habitat) flanked on both sides by cliffs (Wave snail habitat), resulting in two Crab-Wave contact zones per island. The islands were Ramsö (“CZA”, N 58°49’27.8”, E 11°03’43.2”) sampled in July 2013, Inre Arsklovet (“CZB”, N 58°50’00.4”, E 11°08’18.7”), Ramsökalv (“CZC”, N 58°50’04.1”, E 11°02’26.8”) and Yttre Arsklovet (“CZD”, N 58°49’51.4”, E 11°08’00.1”) sampled in May and June 2014 (Fig. S1; for further sampling details see Westram et al. (submitted) but note that CZC is unique to this study). Distances between islands ranged from approximately 0.4 km to 5.6 km. *Littorina saxatilis* has direct development without a pelagic larva and the lifetime dispersal was estimated by Westram et al. (2018) to be about 1.5 m.

“Transect” snails (~600 individuals per location) were collected across the entire length of each transect and their exact positions were recorded in three dimensions using a Trimble total station (as in Westram et al. 2018). “Reference” snails (used as mating partners, see below) were sampled at a fifth island (“ANG” in Westram et al. 2018; N 58°52’15.14“, E 11°7’11.88”) in Crab and Wave habitats away from the contact zone (in total 200 individuals from each habitat per test shore). Both reference and transect snails were sexed prior to mating experiments based on observation of the male penis. If no penis was observed, individuals were assumed to be females. If the penis was underdeveloped, individuals were considered sexually immature and excluded from the mating experiments. Dissections of transect snails followed all experiments in order to confirm initial sex determination and check whether females were immature or parasitised. Trials involving immature or parasitised transect individuals, or individuals whose sex had been determined incorrectly, were discarded.

Size was measured for both reference and transect snails as the maximum distance between the top of the apex and the base of the aperture of the shell. Shape was determined only for the transect snails and summarized as the first relative warp from a landmark-based geometric morphometrics analysis, which captures the Crab-Wave axis of variation (Ravinet et al. 2016; Westram et al. 2018). Shell shape of the reference snails was not analyzed but considered typical of the Crab or Wave ecotype since they were sampled in habitats far from contact zones.

In order to find the relationship between mating probability and the recorded traits (size and shape), we tested each of the transect snails in mating trials with snails from the reference site. Each mating trial involved one transect snail and one reference snail of the opposite sex. The use of reference snails allowed us to avoid confounding mating patterns driven by snail size (or other traits) with patterns driven by population of origin. The use of transect snails from throughout the transects provided a wide range of trait values (and trait value combinations between males and females). Reference snail ecotype and transect snail shape (a continuous proxy for ecotype) allowed us to test for ecotype effects on mating pattern.

Mating trials were performed indoors under constant light and at room temperature. Snails were placed foot-down at the bottom of a transparent plastic sphere (80 mm in diameter) one-third filled with sea water. Plastic spheres were rinsed carefully between trials in order to remove all mucus trails from the previous test. Each transect snail was included in four different trials (on different days) so that it was paired twice with a random Crab reference snail, and twice with a random Wave reference snail. Time of day and ordering effects were avoided using a balanced experimental design. Each mating trial (transect-reference pair) was unique (i.e., involved a pair of snails that was not brought together again) and it was monitored for two hours during which male mounting activity was recorded. Upon encountering another snail, males can crawl onto and around the shell of the other individual until arriving at a characteristic mounting position on the right-hand side of the partner’s shell, inserting the penis under the shell and exploring the mounted snail’s sex. If it is another male, the mating attempt is interrupted, while if it is a female, mating may continue (Saur 1990). Male mounting position is a reliable proxy for a copulation attempt in *L. saxatilis* (Hollander et al. 2005). In addition, a positive correlation between mounting duration and the probability that the female received sperm has been observed in other littorinid species (Hollander et al. 2018). If either the transect or reference snail was inactive throughout the two-hour trial, this trial was excluded from analysis. In the analyses presented here, we considered only whether or not a mating occurred in each trial. A positive outcome was recorded if the male was in the mounting position for more than 1 minute (Saur 1990).

### DATA ANALYSIS

For each mating pair, we had information about whether a mating event was observed or not, the island where the transect snail was collected (CZA, CZB, CZC or CZD), transect snail shape, the ecotype of the reference snail (Crab, Wave), the sex of the transect snail (and therefore of the reference snail) and the sizes of the two snails, which were used to calculate the ratio between the female and male size for each mating pair.

Previous work suggests that the size of the male relative to the female size is the primary determinant of mating, given an encounter (Conde-Padín et al. 2008). We began by checking whether our observations were consistent with this result by fitting generalized linear models to our data. Using the function glm()in R version 3.5.0 (R Core Team 2018) and treating mating as a binary response, we searched for the best models using all possible combinations of seven variables (ln female size], female-male size ratio expressed as {ln female size] - ln male size]}, size ratio squared, size ratio cubed, ecotype of the reference snail, shape of the transect snail and island where the transect snails were collected) and their two-way interactions with the exception of interactions between size variables. The square of the size ratio was expected to account for most of the variance because alone it would generate a decrease in mating probability on either side of a size ratio of zero (i.e., equal size male and female). Any shift in the optimum size ratio away from zero would cause a size ratio effect to be added to the model, as would any asymmetry in the mating probability. The best model, with the lowest Akaike information criterion (AIC = 4251), included effects of size ratio squared and various interaction terms, including two-way interactions between ln(female size) and island, and between size ratio and both shape and island, although their effects were relatively small. Multiple models with similar AIC values consistently included size ratio effects, with the square of size ratio being the strongest effect, but varied in the other factors that entered the model (Table S1, S2).

We then fitted a model to the observed data in order to describe the relationship between mating probability and the ratio of female to male size. We selected a function to account for the decline in probability away from an optimum size ratio (the effect of the square of size ratio in the GLM). This model allowed us to estimate parameters for the mating pattern that we then applied to size distributions in nature to infer the assortative mating and sexual selection generated by the mating pattern. The parameter estimates were also used to simulate the barrier effects of size-assortative mating and sexual selection (see below, CLINE SIMULATIONS). Initial trials showed that a symmetrical Gaussian model that is commonly used to describe sexual selection and assortative mating (Lande 1981; Gavrilets 2004) could not account for our observations because the mating probability declines asymmetrically, more rapidly for males larger than females than for males smaller than females. Therefore, the binary outcome of the mating experiment (mated or non-mated pair) was fitted using logistic regression to a skew normal function of the size ratio. Specifically, we expressed the probability of mating (p_i_) of the *i*-th mating pair as follows:

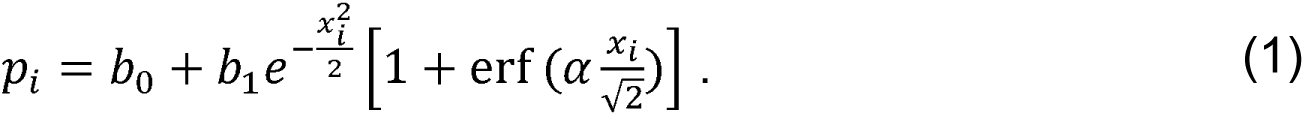

Here, 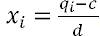, with *q_i_* denoting the observed size ratio on natural logarithm scale, erf is the error function (Glaisher 1871), and *b*_0_, *b*_1_, *c*, *d* and a are (unknown) model parameters (see below). The ‘error function’ provides for an asymmetrical departure from the Gaussian function. For a symmetric model, the probability of mating would be highest for a size ratio of c (called ‘preference’ by Kopp et al. 2018). However, in an asymmetric (skew normal) model, the position of the maximum (the ‘optimal size ratio’, OR) also depends on the parameter *α*, which controls the amount of skew (Fig. 1). The OR was estimated by taking the first derivative of Eq. (1) using Wolfram|Alpha (access October 19, 2018) and finding its root using the function uniroot()in R version 3.5.0 (R Core Team 2018). The rate of decline in the probability of mating away from the OR is given by the parameter *d* (called ‘choosiness’ by Kopp et al. 2018; Fig. 1) and also on *α*. Here we refer to *d* as ‘ratio dependence’ and *c* as ‘center’ to avoid any implication that one or the other sex is making a choice. Finally, parameters *b*_0_ and *b*_1_ are scaling parameters proportional to the overall minimum and maximum proportion of trials in which mating occurred: we call them the ‘mating baseline’ and the ‘mating rate’, respectively (Fig. 1).

**Figure 1.**
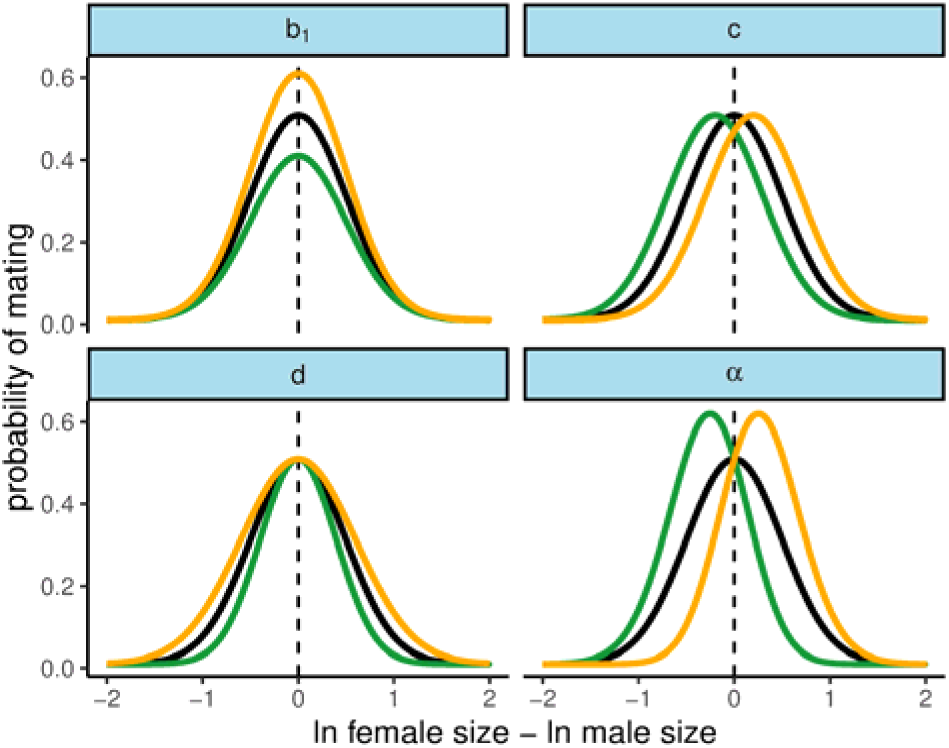
Effects of the parameters on the predicted mating probability. The relationship between probability of mating (y axis) and size ratio (x axis) is determined by five parameters (*b*_0_, *b*_1_, *c*, *d* and *α*). Parameter *b*_0_ is expected to have a low value in all cases and is set here to 0.01. Black lines in all panels have all parameters at the centers of the prior ranges used in model fitting with OR marked by dashed lines. Orange and green lines show the effect of increasing and decreasing a parameter by 20% of its prior range, respectively. Top-left panel – mating rate; *b*_1_, top-right panel - center; *c*, bottom-left – ratio dependence; *d*, and bottom-right - skewness; *α*.

Model fitting was performed in Stan (Carpenter et al. 2017), a probabilistic programming language that adopts full Bayesian statistical inference with Markov Chain Monte Carlo (MCMC) sampling, implemented using the R package ‘rstan’ (Stan Development Team 2018). The space of the parameters was defined using uniform priors that were bounded according to biologically-reasonable limits (0 to 1 for *b*_0_ and *b*_1_; -10 to 10 for *c* and *α*; 0 to 10 for *d*). The sampling algorithm was set to 8000 iterations and it was repeated four times in parallel. The first 2000 iterations of each of the four chains were not used for the posterior inference as these initial values might confound the posterior mean calculations. The rest of the arguments were left at the default settings.

Our initial data exploration using generalized linear models (see above) suggested that the relationship between mating probability and size ratio might vary according to island and ecotype (or snail shape). Furthermore, although the unit of replication was the transect-reference pair, it remained to be tested whether individuals differences in shape between transect snails could have explained part of the variation in mating probability. We tested the impact of these variables by fitting hierarchical models in Stan. In these models, we replaced one or more of the parameters in Eq. (1) by a ‘hyperparameter’ that was a function of the island from which the transect snail was sampled, the transect snail shape, the reference snail ecotype and the sex of the transect snail (Supplementary Information: HIERARCHICAL MODELS; Supplementary Information: MODEL COMPARISONS).

### Mating Pattern Consequences in the Contact Zone

The parameters of the mating pattern were estimated from the observations in the mating experiment, which was designed to investigate the probabilities of mating given encounters between snails with a wide range of sizes and shapes. The implications of this mating pattern for assortative mating and sexual selection in nature depend on the sizes of snails that actually encounter one another. In turn, this depends on how the distributions of male and female size change across the contact zones. It may also depend on dispersal, which determines the spatial scale over which individuals can choose their mates (Rolán-Alvarez et al. 2015). Therefore, we predicted mating of *L. saxatilis* in natural conditions, using the parameters of the skew normal function estimated through Bayesian inference (see above), in order to infer the resulting strengths of assortative mating and sexual selection in our transects.

To obtain the means and variances of male and female size distributions at each point in each transect, we fitted clines to the observed ln(size) data. We estimated cline centers and widths, Crab and Wave ecotype means and the change in variance across the transect by maximum likelihood (‘bbmle’ package in R, function mle2(), Bolker and Team 2017) using equations from Derryberry et al. (2014) and R scripts adapted from Westram et al. (2018) to fit clines at both Crab-Wave contacts simultaneously. Clines were fitted for each island separately using the shell sizes of the transect snails grouped by sex and the position on the shore where they were sampled (on a one-dimensional transect, see Westram et al. submitted) (Table S5).

Mating predictions were run for each of the four islands separately. Each run consisted of repeated sampling of female and male sizes from the fitted phenotypic cline, at multiple positions from one end to the other of the transects. The positions were the island-specific cline centers and a series of equally-distributed distances from the centers for a total of 37 positions in CZA, 26 in CZB, 17 in CZC and 27 in CZD. The positions were separated by a spatial interval of 10 m to ensure sufficient coverage of the contact zone where we expected the size distributions, and thus the intensity of assortative mating and sexual selection, to vary. We assumed that the formation of female-male pairs was constrained to males close to the focal females and that female reproductive success was independent of the number of matings due to their highly promiscuous behavior and capacity for sperm storage in the wild (Panova et al. 2010; Johannesson et al. 2016; but see Ng et al. 2019 who assumed that female fitness increases with number of matings).

At each transect position, T_f_, sizes for 1000 females were drawn randomly from a normal distribution with the mean and standard deviation (SD) predicted for that position on the fitted cline. For each female, we drew a male position T_m_ = T_f_ + ξ, where ξ is a random number from a normal distribution with mean 0, and standard deviation σ = 1 m. We then drew a size for that male using the mean and standard deviation of male size from the cline fit for position T_m_ and determined the probability that an encounter between this pair of individuals would lead to a mating using their size ratio and the skew-normal distribution with our estimated parameters. Whether or not a mating occurred was then determined by a random draw from the Bernoulli distribution with this probability of mating. If no mating occurred, a new male was drawn and the process was repeated until the female mated. We recorded the sizes of males and females in each encounter and the mating outcome. This pipeline was replicated ten times at each position along the transect to obtain reliable estimates of assortative mating and sexual selection.

From the resulting data we extracted the strengths of assortative mating and of sexual selection on males and averaged across the ten runs at each cline position on each island. Assortment was measured as the Pearson correlation coefficient of ln(size) between males and females in mated pairs, while sexual selection was estimated as (i) the difference in mean ln(size) of mated males compared to mated plus non-mated males (directional component) and (ii) the difference between the variance of ln(size) for mated males and for all males (stabilizing component).

### CLINE SIMULATIONS

The observed mating pattern reflected the extent of assortative mating, displacement of the optimum size ratio from zero (i.e., from equal male and female sizes) and asymmetry of the mating function. To understand how these effects contribute to the barrier to gene flow between ecotypes, we performed individual-based computer simulations for the evolution of a cline across a contact zone comparing five models that sequentially add these effects. We take the width of the trait cline as a measure of barrier strength because it is expected to reflect the impact of the barrier on gene flow (Barton and Gale 1993): a narrow cline implies a strong barrier. In each model, the habitat consisted of 400 patches arranged linearly, each with 100 diploid individuals (50 males and 50 females). Consecutive patches were assumed to be 1 m apart. Generations were discrete and non-overlapping. The lifecycle was modelled in the order: dispersal, recombination and mating, locally in each patch, then natural selection. In the model, dispersal distance was Gaussian distributed with mean zero and standard deviation σ = 1.5 (in line with the estimate in Westram et al. 2018). We assumed that the trait under selection (i.e., the size of individuals on a natural logarithmic scale) had an optimum (*θ_j_*) that changed abruptly at the center of the habitat (between the patches 200 and 201), so that *θ_j_* = 2 for patches *j* = 1,2, …, 200, and *θ_j_* = −2 for patches *j* = 201,202, … ,400. Since size is typically a polygenic trait (Houle 1992), the modelled trait under selection was assumed to be underlain by a large number of loci (but not too large, for computational efficiency), i.e. we assumed a set of L = 40 loci in females, and a separate set of L = 40 loci in males (but we traced the evolution at all 80 loci in all individuals). Separate sets of loci underlying the trait under selection were used since this is the simplest architecture that allows sexual dimorphism to evolve. All loci were assumed to recombine freely. Each locus had additive alleles of effect size 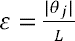 or −*ε*, so that, due to diploidy, overshooting of the local trait optimum was possible. Mating was implemented according to five different models, one being random mating, and the remaining four being different versions of the mating pattern based on the trait that was also under natural selection (see below). In each model, we assumed that every female produced a large (and the same) number of offspring (i.e., 100), so that there was no sexual selection on females. By contrast, males could have different contributions to the pool of offspring, as a result of the mating model applied. After reproduction, the adults died, and the pool of offspring in each patch was randomly divided into 50% males and 50% females. To keep the population size constant, we then applied natural selection so that only 50 females and 50 males survived in each patch. The fitness *W_k,j_* of an individual *k* in patch *j* depended on the distance of the individual’s trait value *Z_k,j_* from the optimum *θ_j_* in the patch according to

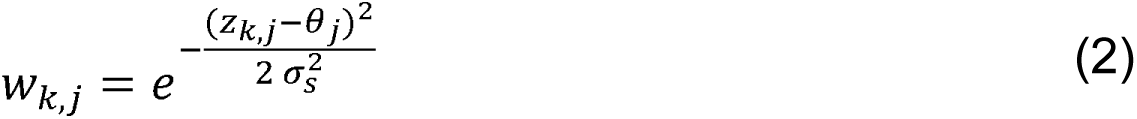

Here, *σ_S_* as is the inverse of the strength of natural selection. We chose it so that an individual that was perfectly adapted to one habitat end had a fitness equal to 0.7 in the other habitat end, and vice versa. This corresponds to a selective disadvantage of 0.3 (chosen on the basis that selection on size is expected to be strong, but not as strong as the total selection against a snail of one ecotype in the wrong habitat, cf. Westram et al. 2018).

As mentioned above, we simulated five different mating models. Random mating (hereafter *RM* model) provided a baseline against which we compared the observed mating pattern that includes assortative mating with a skewed mating probability according to Eq. (1) with our estimated parameters (hereafter *AS* model). The *AS* model contains effects of assortative mating, directional sexual selection due to displacement of the optimum and directional sexual selection due to the asymmetric mating probability. We separated these effects by adding three other models. First, assortative mating without directional sexual selection was simulated using a symmetric mating probability with mean 0, and standard deviation equal to our empirical estimate of d (hereafter *SimM0* model). Then, we simulated two models with symmetric mating probability but a shifted optimum, either equal to the optimum of Eq. (1) (hereafter *SimOR* model) or to the mean of Eq. (1), which is shifted further from zero than the optimum as a result of the skew (hereafter *SimMR* model) (both with standard deviation equal to d). These models allow us to test the effect of assortative mating alone (*SimM0*), assortative mating combined with a shift in the optimum trait ratio corresponding to either the observed mean (*SimMR*) or the observed optimum (*SimOR*) but without asymmetry, or the full observed mating model including skew (*AS*). Model *AS* always generates both stabilizing and directional sexual selection on males. Models *SimXX*, where *XX is either MR, OR* or *M0*, generate only stabilizing sexual selection whenever the sexual dimorphism present in a population matches the optimum of the mating function and do not generate any sexual selection when the distribution of male mating probability fully matches the distribution of males in the population. Note that the mating pattern did not evolve in any of our models. Rather, it was fixed both in space and time. This was because empirical data did not show any significant differences in the mating pattern between ecotypes or islands (see Results) and other work suggests that mating patterns are similar in other littorinids (Ng et al. 2019). Therefore, our model did not account for any genetic variation in the mating pattern and it was not necessary to specify its genetic basis in order to investigate its expected effect on the current barrier to gene flow.

Each simulation was initialized so that alleles of effect size ε were fixed in patches *j* = 1,2, …,200 at all loci (and −*ε* in patches *j* = 201,202, …,400) such that all phenotypes were either 2*θ* or −2*θ* at the start of the simulations, and for this reason mutation was not required. We then ran each simulation under the random-mating model until approximately a steady state was reached, that is, for 10,000 generations (burn-in period). We performed 200 independent realizations for this burn-in period, and we used the results from the last generation of the burn-in period as initial conditions for the simulations with assortative mating (same initial conditions for each of the four models; see above). We then ran each model with assortative mating for an additional 5,000 generations, during which the population reached approximately a steady state.

For the burn-in period (random mating), and for the runs with assortative mating, we collected simulation results from the final generation simulated in each case. We estimated a hybrid index across all loci (i.e., both those influencing male and those influencing female size). The hybrid index was the relative frequency of alleles with effect sizes *ε* averaged over all loci. It was expected to run from 0.75 in patch 1 to 0.25 in patch 400 in the steady state with random mating such that phenotypes run from 2 to −2 and it was calculated in each patch, separately for males, females and all individuals. We then fitted clines to the hybrid index using equations from Derryberry et al. (2014) including symmetric, asymmetric, and tailed clines, with one or three independent variances, and R scripts adapted from Westram et al. (2018). In addition, we fitted the spatial pattern of the hybrid index obtained in our simulations to a constant value, independent of the spatial position (which is an expected pattern under neutral evolution), to check whether a clinal pattern explains our hybrid-index data better than a neutral-evolution model (using AIC). This was indeed the case (see Results). In the great majority of cases, the best fit was achieved by a symmetric cline model with left and right tails and three independent variances (not shown). For each realization, the maximum-likelihood values for the estimated cline centers, widths (i.e., 1/slope at the cline center), and hybrid index at the habitat ends were saved for comparison between the different models. Specifically, we approximated the inverse strength of the reproductive barrier in a given model by the estimated cline width for the best fitting model (scaled by the difference of the hybrid index between the habitat ends). Thereafter, we compared the strength of the reproductive barriers established in the different models by investigating the distributions of the estimated cline widths obtained in the different models in 200 independent realizations.

## Results

The raw number of mating trials (all islands included) was 7594 and, after the filtering steps, 4330 trials were used for the downstream analysis. The excluded observations contained 530 mating pairs where the sex of the transect snails was misidentified, 968 where stage of the transect snail was juvenile, 292 with parasitised transect snails, 1286 where one or both snails were inactive throughout the trial, 70 transect snails without spatial information and 118 mating pairs with missing shell sizes.

### Mating Pattern in the Laboratory Trials

Size-assortative mating acts as a barrier to gene flow when the probability of mating between two populations of different sizes is reduced. To investigate this barrier effect, the first step is to quantify how the probability of mating varies with respect to female and male size distributions. The mating model, Eq. (1), was built for this objective and it was fitted to the data from all four islands combined (Fig. 2). The probability of mating followed a right-skewed distribution with maximum displaced from the center of the distribution towards pairs where the female was 1.31 times larger than the male and falling rapidly for pairs with other size ratios (Table 1; Fig. 2). As the size ratio between the sexes increased/decreased, the mating function approached a probability close to zero within the range of observed size ratios for males larger than females but not for males smaller than females (Table 1; Fig. 2). To give an example of what these values mean in practice, a female of 12.5 mm had the highest probability (0.56) to mate with a male of 9.5 mm (~25% smaller, optimal ratio = 0.27). The same female would mate with a 5.2 mm male with probability 0.33 or with a 17.4 mm male with probability 0.25, despite their size ratios [on ln scale; 0.87 and −0.33, respectively; ln(female size) - ln(male size)] being equidistant from the optimal ratio (OR). With such an asymmetric pattern, smaller males have a mating advantage over larger males even when sexual size dimorphism is such that the average size ratio corresponds to the optimum ratio for mating.

**Table 1.** Parameter estimates for the non-hierarchical model. The summary statistics are mean, standard deviation (SD), lower bound of 95% CIs (2.5%), upper bound of 95% CIs (97.5%). Optimum size ratio (OR) and the mating probability at this ratio were derived from the fitted parameters, with confidence intervals derived from the MCMC chain.

| Parameter | Mean | SD | 2.5% | 97.5% |
| --- | --- | --- | --- | --- |
| $b_0$ | 0.01 | 0.01 | 0.00 | 0.02 |
| $b_1$ | 0.40 | 0.02 | 0.36 | 0.46 |
| $c$ | -0.17 | 0.04 | -0.23 | -0.09 |
| $d$ | 0.85 | 0.06 | 0.74 | 0.97 |
| $\alpha$ | 2.33 | 0.39 | 1.61 | 3.15 |
| OR | 0.27 | 0.03 | 0.21 | 0.32 |
| Mating probability | 0.56 |  | 0.53 | 0.59 |
| at OR |  |  |  |  |

**Figure 2.**
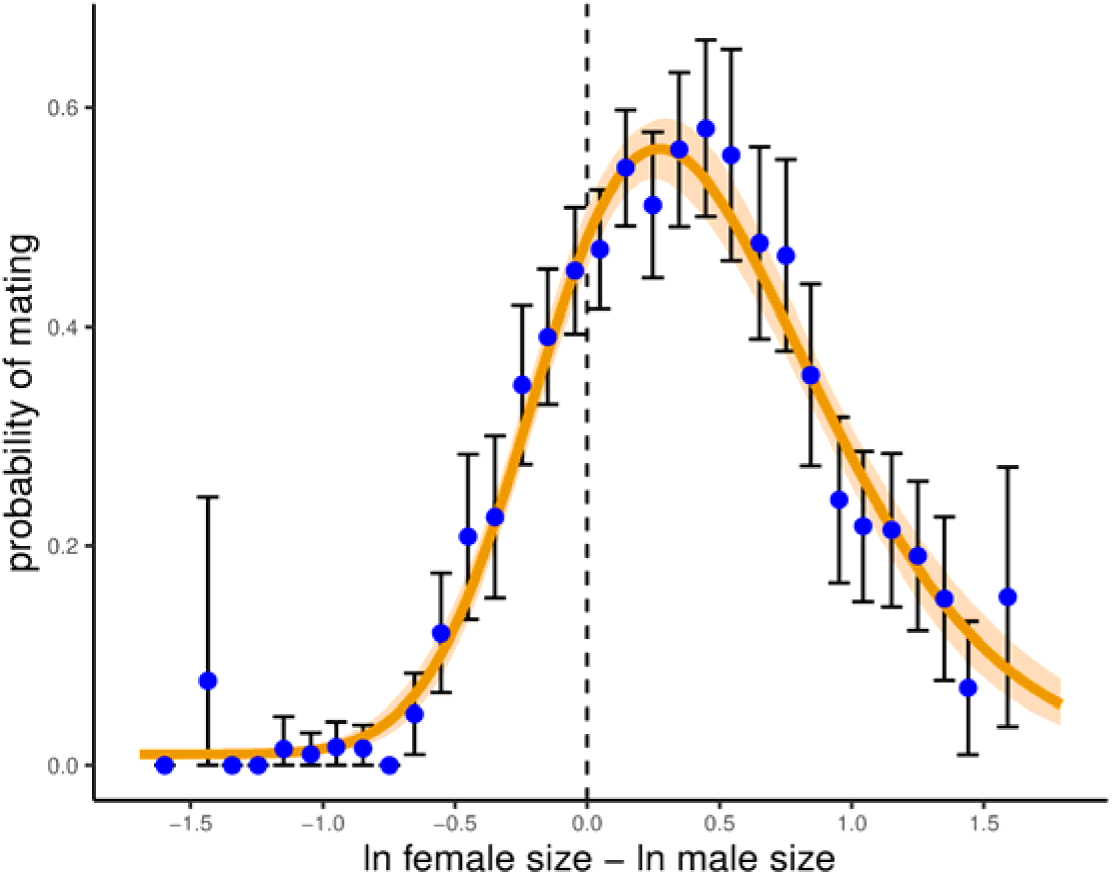
The mating pattern across all islands, fitted by the non-hierarchical model, followed a right-skewed distribution. Fitted curve and 95% CIs in orange are superimposed on the observed proportions of matings (Blue dots - proportions of trials resulting in mating for size ratio bins. Black error bars – 2.5^th^ and 97.5^th^ percentiles).

Fitting of hierarchical models showed some statistically significant but small improvements in the explanation of mating pattern: mating rate (parameter *b*_1_) varied among islands and between sexes of the transect snails, and the center parameter (c) varied slightly between islands and between reference ecotypes (Supplementary Information: HIERARCHICAL MODELS; Supplementary Information: MODEL COMPARISONS). Given the small effect sizes, especially for difference in pattern as opposed to rate of mating, in the following predictions and computer simulations we used the non-hierarchical model (i.e., the model where the mating pattern was considered invariant within and among ecotypes and islands).

### Assortative Mating and Sexual Selection

Clines in male and female size were observed on all four islands with centers close to habitat boundaries (Fig. 3; Fig. S4). In all cases, sexual size dimorphism was greater in Wave snails than in Crab snails, the variance in ln(size) was also greater in Wave snails and the variance increased in the centers of the clines (Fig. 3; Fig. S4; Table S5).

**Figure 3.**
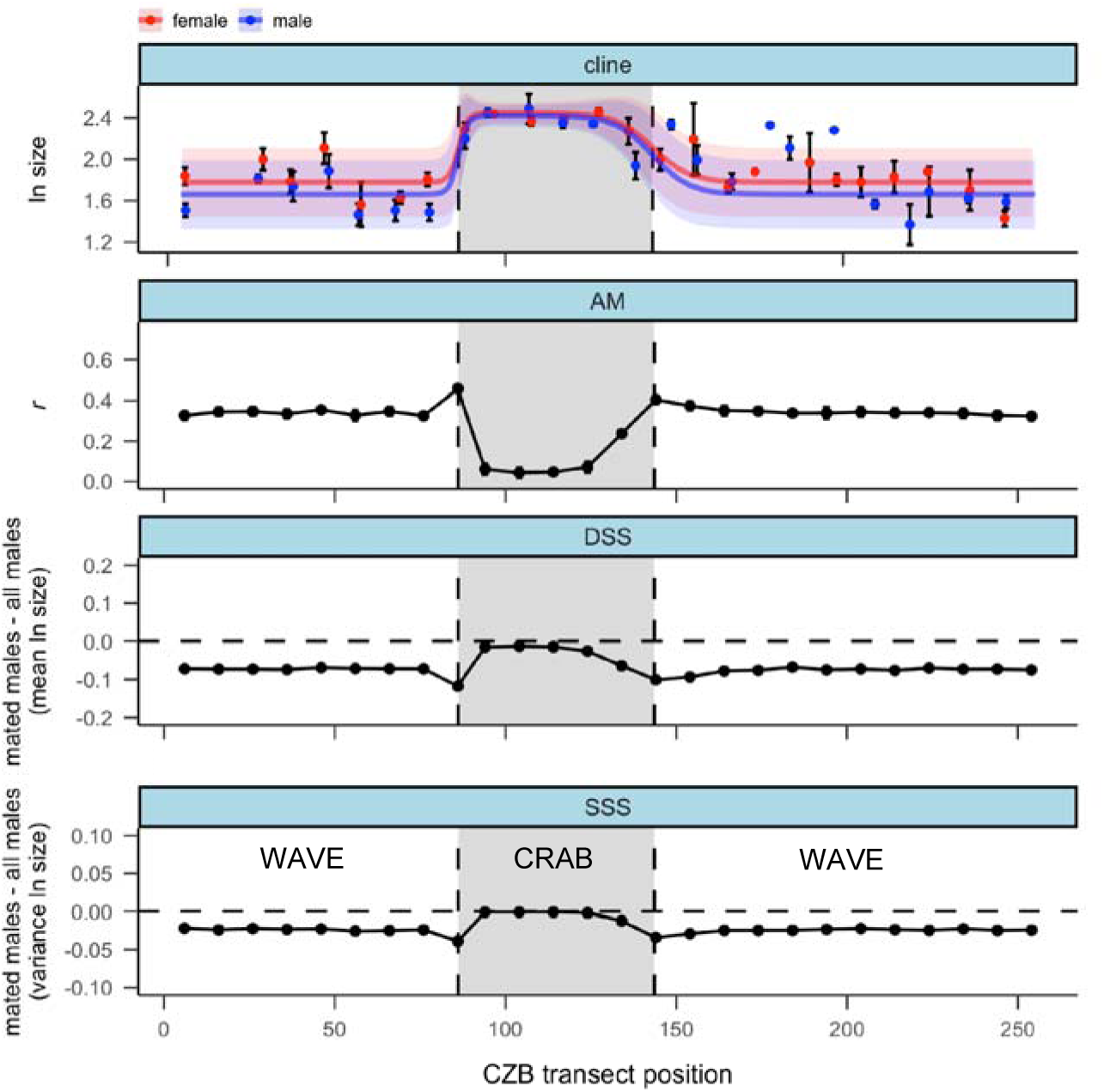
Predicted assortative mating and sexual selection (CZB transect as an example: for the other three transects, see Fig. S4). Habitat boundaries are marked by black vertical dashed lines, the crab habitat is the region inside (grey fill) and the wave habitat is outside (white fill) the two dashed lines. Cline facet: ln(size) of transect snails in bins (dots with 95% CIs) and fitted clines (solid lines ± SD) for females (in red) and males (in blue). AM facet: strength of assortative mating measured as the Pearson correlation coefficient (r) between female and male ln(size) of mated pairs. DSS facet: directional component of sexual selection measured as the difference in mean ln(size) of mated males compared to mated plus non-mated males. The black horizontal dashed line indicates where this component is absent. SSS facet: stabilizing component of sexual selection calculated as the difference in variance between mated male ln(size) and mated plus non-mated male ln(size). The black horizontal dashed line indicates where this component is absent.

After generating virtual mating encounters using a custom script, we computed, for each position along the transect on a specific island, the correlation (Pearson’s *r*) between female and male ln(sizes) in the virtual mated pairs (i.e., assortative mating) and the difference in mean and variance of ln(size) of mated males compared to all the males that were generated at that particular transect position (i.e., sexual selection). Positive size-assortative mating was predicted for all transect positions in all four Swedish islands. Predicted assortment was strongest at the centers of the clines where the size variance was greatest, intermediate in the wave habitat and weakest in the crab habitat where the size variance was smallest (Fig. 3; Fig. S4).

Sexual selection was predicted to favor smaller males, and lower variance in male size in all cases (Fig. 3; Fig. S4). However, sexual selection was also predicted to vary along the transects of the four islands in line with the size variance and difference between female and male sizes. In some cases, the predicted effects were very small. Specifically, the directional component (DSS, the difference in mean between mated and all males) was most negative at the centers of the contact zones (where the variance of ln(size) of males was highest), intermediate in the wave habitat (where variance in ln(size) was intermediate) and close to zero in the crab habitat (where the variance in ln(size) was lowest). The stabilizing component of sexual selection (SSS, the difference in variance between mated and all males) showed a similar pattern to the directional component.

### Barrier to Gene Flow

In all five models we simulated (see illustrations in Fig. 4a, e, I, m, q) we found that, at the end of the simulations, the average phenotype of females at the two habitat ends matched their corresponding optimal phenotypes (Fig. 4b, f, j, n, r, solid red lines). For males this was only true under the random mating model and under the *SimM0* model (Gaussian mating probability with optimum at zero, i.e., with the maximum mating probability for equal-sized males and females; see Fig. 4b and Fig. 4r, where the blue solid line overlaps with the red solid line). In the remaining three models, in each patch males attained on average smaller phenotype values than females (Fig. 4f, j, n). For symmetric mating functions (Fig. 4j, n) the difference between the optimal phenotype and the average phenotype attained by males at either habitat end approached the optimum of the corresponding mating function, indicating that directional sexual selection on males was strong relative to the natural selection implemented in the model. Conversely, when the mating function was asymmetric (Fig. 4f), the difference was slightly larger than the optimum of the function (dashed blue line). This is because the mean of the mating function, Eq. (1), was slightly larger than its optimum due to the asymmetry (compare dashed cyan line to the dashed blue line in Fig. 4e). The difference between the final phenotype of males and their optimal phenotype under natural selection alone was slightly larger than the mean of the mating function (blue solid line is between dashed blue and dashed cyan line in Fig. 4f). This is because natural selection (that acts after mating) favors males with the phenotype closer to the optimum, and the relative contribution to the overall fitness of males further away from the optimum was disproportionate in comparison to the contribution of males closer to the optimum. This made the component of natural selection acting on males effectively stronger in the case of the asymmetric mating function (*AS* model, Fig. 4f) than in the case of a symmetric mating function with the optimum equal to the mean of the asymmetric function (*SimMR* model, Fig. 4j).

**Figure 4.**
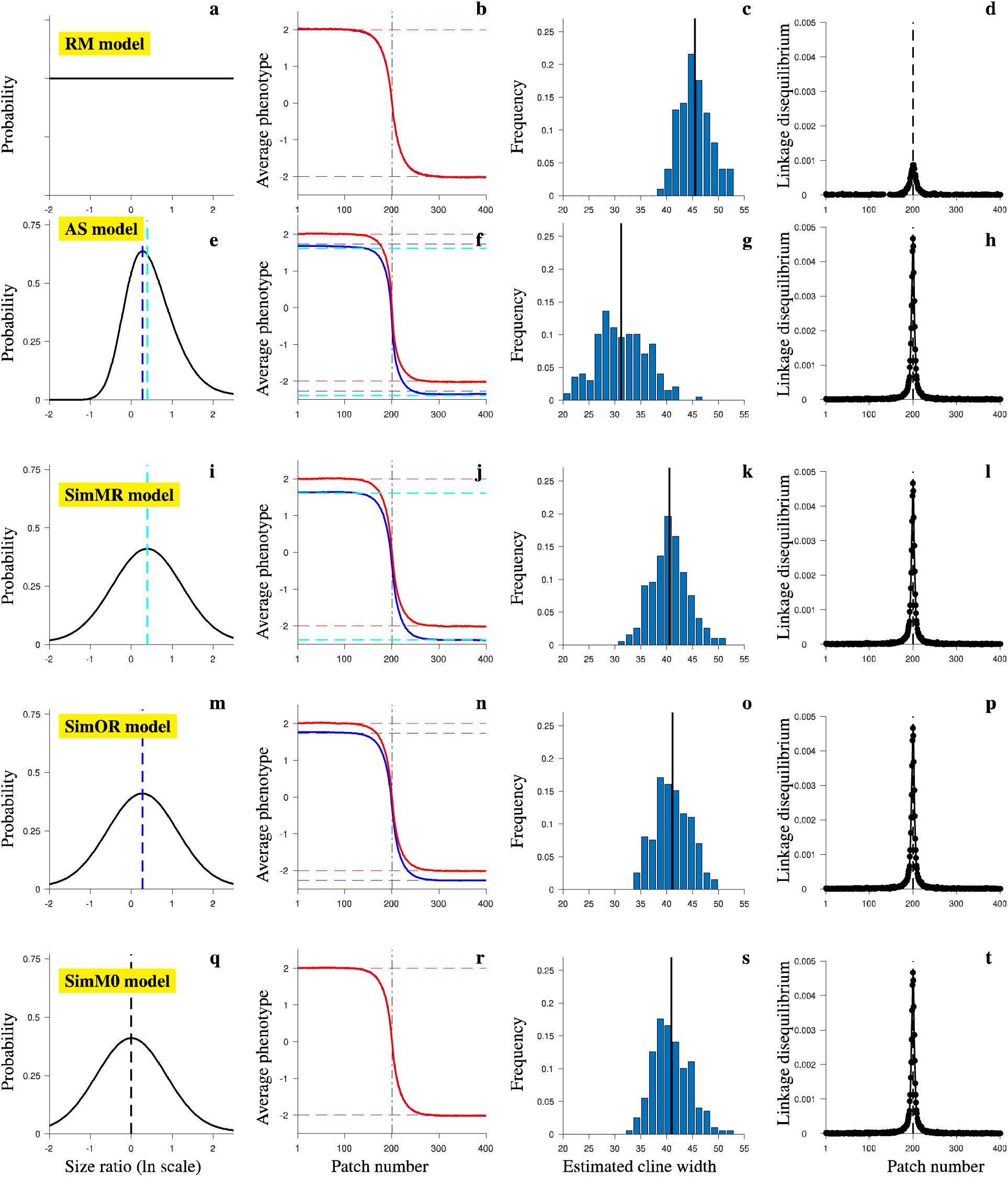
Mating models simulated and the simulation results showing that the strongest barrier (i.e., narrowest clines) is generated by the observed (*AS*) mating pattern. First column: probability of mating as a function of the size ratio between males and females on ln scale in *RM* model (a), *AS* model (e), *SimMR* model (i), *SimOR* model (m), and *SimM0* model (q). In a), the mating probability is independent of the size ratio, and the scales on the x- and y-axis are chosen arbitrarily for illustrative purposes. Blue dashed lines denote the optimal ratio (OR) in (e, m). Cyan dashed lines denote the mean ratio (MR) in (e, i). Black dashed line denotes MR in (q). Note that MR and OR are equal in (i, m, q). Second column: average phenotype at the end of the simulations as a function of the patch number. Solid lines show the phenotypes of females (red), or males (blue). Note that blue and red lines overlap in (b) and (r). Dashed lines show the optimal phenotype at the two habitat ends (θ_1_ and *θ*_400_ = −*θ*_1_; red), optimal phenotypes at the two habitat ends minus OR (blue; f, n), and optimal phenotypes at the two habitat ends minus MR (cyan; f, j). Vertical dash-dotted line shows the position of the environmental transition. Third column: distribution of estimated cline widths for the hybrid index, considering all individuals at the end of the simulations. Vertical lines show the mean values. Fourth column: average linkage disequilibrium as a function of the patch number at the end of the simulations. Dashed lines denote the position of the environmental transition. 200 independent realizations of each model.

We computed a hybrid index (HI) in each patch (proportion of alleles with positive effect sizes averaged over all 80 loci), and for each realization of the different models we fitted clines. The spatial pattern of HI was best explained by a symmetric cline model in all cases (not shown). As a proxy for the overall inverse strength of the reproductive barrier in each case, we measured the corresponding cline widths (Fig. 4, third column). The cline widths for the model with asymmetric mating function (*AS* model) were significantly smaller than cline widths for the random-mating model (compare Fig. 4c, and Fig. 4g, as well as the first and second rows in Fig. S5): a width of less than 40 patches was found in only about 3% of clines obtained under the random mating (*RM*) model, but in 97% of clines under the *AS* model. Thus, the barrier was statistically significantly stronger than in the random-mating case: on average, the cline width in the *AS* model was smaller by about 31% than in the *RM* model, and by about 23-24% compared to the *SimXX* models, with slight differences between the individual symmetrical models that differ only in the position of the peak mating probability. In other words, the barrier (1/width) in the *AS* model was stronger by about 46% than in the *RM* model (and about 30-32% stronger than in the *SimXX* models). We found that assortative mating also increased the barrier strength in comparison to that established under the *RM* model for the remaining three models of assortment (compare solid vertical lines in Fig. 4k, o, s to the vertical line in Fig. 4c; see also Fig. S5), but the difference to the *RM* model was not as great as in the case of the *AS* model. Among the three symmetric mating models we simulated, the barrier strength was strongest for the *SimMR* (peak at the observed mean; Fig. 4k; Fig. S5, third row), and slightly weaker in the *SimOR* model (peak at the observed optimum; Fig. 4o; Fig. S5, fourth row), and in the *SimM0* model (peak at zero; Fig. 4s; Fig. S5, last row). This was because any deviation of the optimum of the mating function from zero introduced a sexual selection component on males (always for lower trait values) taking their phenotype away from the natural selection optimum. The component of natural selection was, therefore, stronger when sexual selection was at work (recall that natural selection pushed the males towards the same phenotype optimum as that for females). However, the differences between the three symmetric mating models were subtle.

There were no significant differences in the distribution of the estimated cline widths between HI clines for males only (first column in Fig. S5), for females only (second column in Fig. S5), or for all individuals (third column in Fig. S5).

In all cases, assortative mating (and sexual selection on males) introduced stronger stabilizing selection on males than on females, resulting in a narrower distribution of phenotypes of males than of females (Fig. S6, second to last row). In the random-mating model, by contrast, the two distributions were indistinguishable, as expected (Fig. S6, first row).

Finally, in all cases with non-random mating the average linkage disequilibrium between pairs of loci was strengthened by a factor of about five (Fig. 4d, h, l, p, t).

## Discussion

Single traits with multiple barrier effects potentially make a strong contribution to the formation of new species as they can overcome the opposition of gene flow and recombination during the build-up of reproductive isolation (Servedio et al. 2011; Smadja and Butlin 2011; Kopp et al. 2018). However, the contribution that such a trait makes to the overall barrier to gene flow has not been measured in the appropriate context, i.e., where hybridization generates intermediate phenotypes. Here, we investigated the contribution to reproductive isolation of shell size, a single trait with effects on both ecological and sexual isolation between Crab and Wave ecotypes of *Littorina saxatilis*. Our results confirm previous observations of size-assortative mating in *L. saxatilis*: mated pairs showed a positive correlation with respect to size. However, our quantification of the mating pattern demonstrates that it also generates sexual selection on male size, with a stabilizing component and a directional component due to a shift in the optimum size ratio towards males smaller than females and an asymmetry in the rate of decline in mating probability either side of the optimum. We show that the strength of assortative mating and sexual selection is expected to vary across contact zones as the male and female size distributions change, despite constancy of the mating pattern itself. We then show, by simulation, that when sexual selection is included into the model of mating pattern, it increases the barrier to gene flow even when it is uniform rather than divergent. The barrier strength due to assortative mating alone is clear. This indicates that the directional components of sexual selection (displacement of the optimum size ratio and, especially, asymmetry of the mating function) are more important than assortment alone in the evolution of divergence in *L. saxatilis*.

Assortative mating is widespread across animal taxa (Janicke et al. 2019). In most marine gastropods studied, females and males mate assortatively in relation to size (Ng et al. 2019). It is also common for the optimum size ratio for mating to involve females larger than males and this is true for populations of *L. saxatilis* remote from our Swedish study sites as well as for related species of *Littorina* (Ng et al. 2019). Together with our finding that mating pattern was very similar among islands and between ecotypes, this suggests that the pattern is ancestral and strongly conserved. The reasons for this are unknown but may relate to the physical constraints on internal fertilization imposed by the gastropod shell. Ng et al. (2019) suggest that the mating pattern generates sexual selection for larger female size. However, at least in *L. saxatilis*, female reproduction is not limited by mating (Panova et al. 2010) and so we expect the major effect to be sexual selection for smaller male size. This is likely to result in sexual size dimorphism, which is commonly observed in marine gastropods (Ng et al. 2019). Given the constancy in mating pattern, the extent of dimorphism is expected to depend on the pattern of natural selection on males and females, in terms of both optimum and intensity. Our data show consistently greater size dimorphism in the Wave ecotype than in the Crab ecotype (Fig. 3; Fig. S4). The most likely explanation for this is strong selection for large size imposed on both sexes by crab predation in the Crab environment (Johannesson 1986).

The asymmetry that we observed in the mating pattern has not previously been reported. It contributes strongly to the directional component of sexual selection because for a given distance from the mating optimum, males that were smaller than females mated with higher probability than males that were larger than females. This means that the directional component of sexual selection is not only present when the mating optimum and the mean size ratio differ but also when they are equal. Since sexual selection can contribute to the barrier to gene flow, this is important. Mating functions used in theoretical studies are invariably symmetrical (Kopp et al. 2018), with obvious benefits in terms of simplicity and tractability. However, our results suggest that asymmetric functions should be considered in future theoretical and empirical work. Our quantitative description of the mating pattern in *L. saxatilis* allowed us to simulate its impact on the barrier to gene flow between ecotypes. This simulation used parameters estimated from the field wherever possible but necessarily made some assumptions. For example, we know that there is divergent selection on size and that it is likely to have a polygenic basis (Janson 1983; Westram et al. 2018) but we had to make assumptions about the specifics of the genetics and of the natural selection function. In particular, we made the simplifying assumption that natural selection works equally on males and females, in the absence of evidence to the contrary. The simulation predicts the impact of the mating pattern, if these assumptions are correct, rather than estimating the actual effects. Nevertheless, our simulation results showed clear effects on barrier strength and allow general conclusions to be drawn. The barrier to gene flow was strengthened by the mating pattern observed (assortative mating plus a component of sexual selection on males) in comparison to random mating (as shown by the narrower clines under the *AS* model, compared to the *RM* model; Fig 4). The *SimM0* mating model allowed us to ask how much of this barrier enhancement was due to assortative mating as opposed to directional sexual selection. With this mating pattern, sexual selection was absent or weak and mainly stabilizing, if the male size distribution differed from the distribution of mating probability. Here, there was an increase in barrier strength, but only by about 10% whereas the observed mating pattern (*AS* model) generated an increase by about 46% (based on the inverse of the mean cline width).

We suggest that this difference can be explained as follows. The increase due to assortative mating comes mainly from an increase in linkage disequilibrium (Fig. 4) which causes individual loci underlying size to experience a stronger component of indirect selection (cf. Barton and Bengtsson 1986). By contrast, sexual selection under the *AS* model creates a much greater increase in the total strength of direct stabilizing selection on males because their phenotypic distribution has to reach a compromise between the forces of natural and sexual selection. As the directional component of sexual selection moves the male mean size further from the environmental optimum, the strength of natural selection back towards the optimum increases. At equilibrium, this is balanced by directional sexual selection, resulting in stronger net stabilizing selection. Essentially, males experience two opposing sources of selection leading to a sharper net fitness peak than under natural selection alone. Since the mating effect does not differ between environments, the two fitness peaks are separated by the same phenotypic distance as they would be under natural selection alone. However, the fitness of a Crab male in the wave environment (or vice versa) is more strongly reduced. This stronger overall selection decreases cline width (increases the barrier effect) despite the fact that sexual selection is favoring small males in both habitats.

Under the other mating models that we simulated (*SimMR* and *SimOR*), there was a directional component to sexual selection in the absence of sexual dimorphism but this was largely removed once dimorphism had evolved. As a result, the barrier effect under these models was very similar to the effect under the *SimM0* model. This may be one area where the models fail to capture important features of the natural system: sexual size dimorphism was quite different between Crab and Wave ecotypes in our field data, showing that no single level of dimorphism can resolve the conflict between natural and sexual selection. Nevertheless, it is clear that the sexual selection generated by the mating pattern asymmetry is likely to generate a key component of the overall barrier effect.

Our results broadly agree with Irwin’s (2019) conclusion that assortative mating alone adds rather little to the barrier effect created by natural selection in a hybrid zone. Because we considered a multiple-effect trait, whereas Irwin considered a signal-preference interaction that was separate from the trait under natural selection, we might have expected a stronger effect. However, our simulations are difficult to compare because Irwin considered a simple genetic basis, resulting in discrete phenotypic categories, and mating rules that were not based on observation and do not relate easily to our description of the mating pattern. Our results reinforce the point that isolation indices from mate choice experiments with parental classes, giving values as high as 0.96 in *L. saxatilis* (Johannesson et al. 1995), are a poor guide to the barrier effect of assortative mating in a hybrid zone.

There is broad theoretical agreement that multiple-effect traits favor the evolution of reproductive isolation (Kirkpatrick & Ravigné 2002; Servedio et al. 2011; Smadja and Butlin 2011; Kopp et al. 2018). However, mating patterns can also impede divergence (Servedio 2011; Servedio and Hermisson 2019). In the *L. saxatilis* case, a preexisting mating pattern of the sort that we now observe (constancy of the mating pattern across islands and ecotypes) would have had contrasting effects on the origin of the Crab and Wave ecotypes: it would have opposed initial divergence but it would have enhanced the barrier effect created by divergence in size. Our current simulations do not address this early phase of ecotype formation which was instead explored by Sadedin et. al (2009). Similarly, we have not addressed the possible ongoing evolution of the mating pattern because we assumed constancy in time and no genetic variation for the mating pattern. We find no difference in mating pattern between ecotypes. Stronger assortment near to cline centers is due to segregating variation in size, rather than any change in mating patterns. The direction of sexual selection is the same across the habitat boundary. Therefore, there is nothing in our data to suggest ongoing evolution of the mating pattern. Reinforcement is unlikely in Swedish *L. saxatilis* because hybrid zones affect only a small proportion of the population and they are subject to strong gene flow from parental populations, which are not conditions likely to generate a response to reinforcing selection (Servedio and Noor 2003). Further evolution of the mating pattern may be more likely in Spanish populations where there is more widespread contact (Galindo et al. 2013).

Finally, while we have shown that assortative mating can strengthen the overall barrier to gene flow in the presence of ongoing hybridization, the effect is weak, even for a multiple-effect trait. A component of sexual selection can enhance the barrier effect, even if it is not divergent. For the mating pattern to generate a strong barrier it would have to involve a much more tightly-constrained pattern of mating.

