## Supplementary Information for "Assortative mating, sexual selection and their consequences for gene flow in *Littorina*"

**SAMPLING, PHENOTYPES AND MATING EXPERIMENT**

CZA	CZB

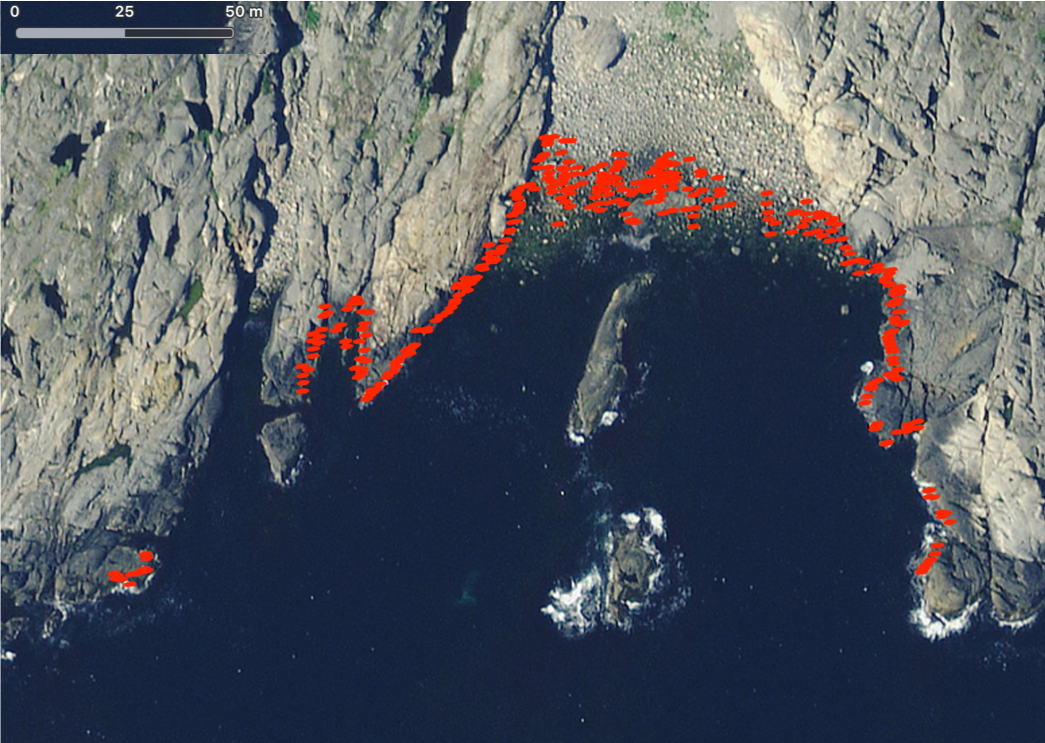

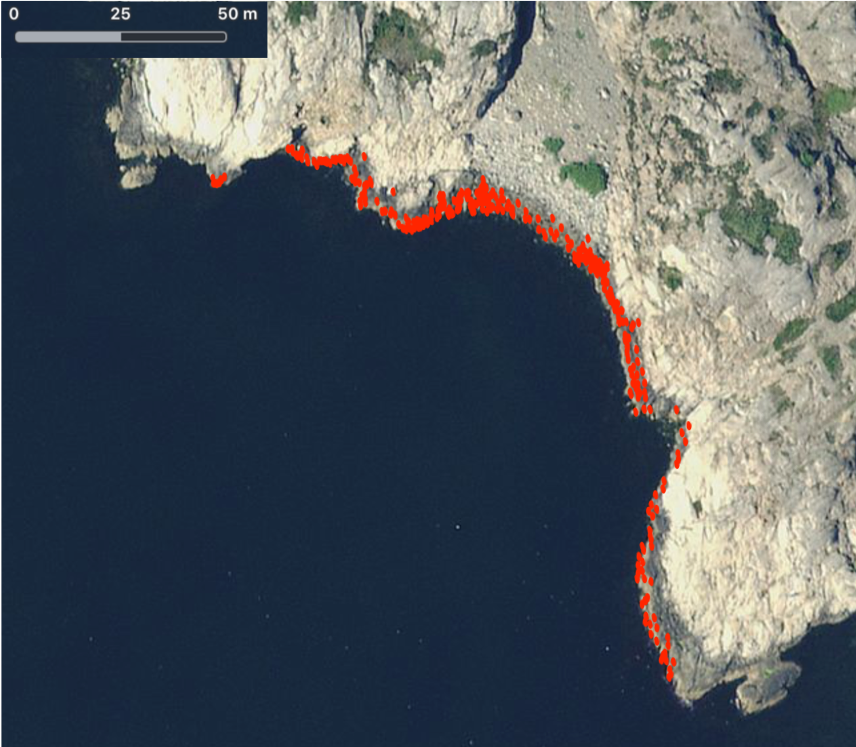

CZC	CZD

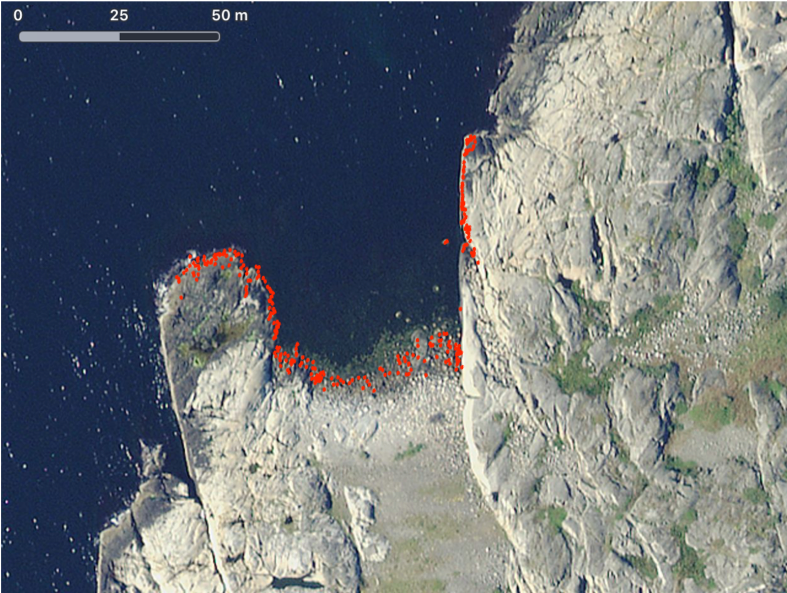

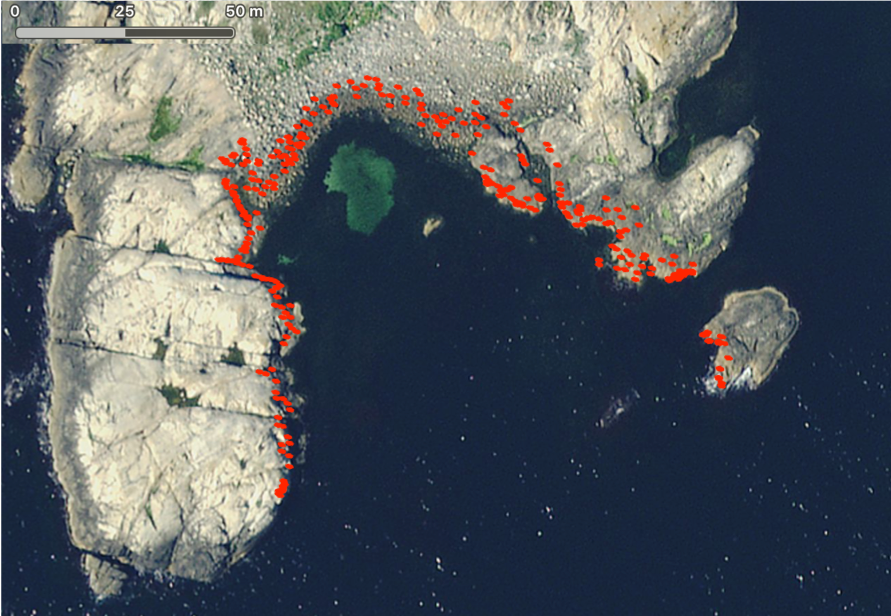

**Figure S1.** Satellite view of the four sample sites (CZA, CZB, CZC, CZD) located on the Swedish west coast. Transect snails were intensively sampled on the four islands (red dots) which included one crab habitat with boulders in between two wave habitats with cliffs, while reference snails for mating trials were all from ANG (for sampling details, see Westram et al. 2018; images from the application Maps © 2012-2018 Apple Inc.).

**Table S1.** Top generalised linear models, with Akaike information criterion ($\text{AIC}$) up to 2 units greater than the best model, among $35442$ models for the probability of mounting. $\text{AIC}$ and formulas of the relationship between the binary dependent variable (‘mountYN’) and the independent variables and two-way interactions.

| **AIC** | **Formula** |
| --- | --- |
| $4251$ | $\text{mountYN}\text{ \textasciitilde size\_ratio2+log\_female:shore+shape:size\_ratio2+shape:ref\_ecotype+size\_ratio:ref\_ecotype+shore:size\_ratio3}$ |
| 4251.2 | $\text{mountYN}\text{ \textasciitilde size\_ratio2+log\_female:shore+shape:ref\_ecotype+size\_ratio:ref\_ecotype+shore:size\_ratio3}$ |
| 4251.2 | $\text{mountYN}\text{ \textasciitilde log\_female+size\_ratio2+log\_female:shore+shape:ref\_ecotype+size\_ratio:ref\_ecotype+shore:size\_ratio3}$ |
| 4251.2 | $\text{mountYN}\text{ \textasciitilde shape+size\_ratio2+log\_female:shore+shape:ref\_ecotype+size\_ratio:ref\_ecotype+shore:size\_ratio3}$ |
| 4251.2 | $\text{mountYN}\text{ \textasciitilde size\_ratio+size\_ratio2+log\_female:shore+shape:ref\_ecotype+size\_ratio:ref\_ecotype+shore:size\_ratio3}$ |
| 4251.2 | $\text{mountYN}\text{ \textasciitilde size\_ratio2+size\_ratio3+log\_female:shore+shape:ref\_ecotype+size\_ratio:ref\_ecotype+shore:size\_ratio3}$ |
| 4251.3 | $\text{mountYN}\text{ \textasciitilde size\_ratio2+log\_female:shore+shape:ref\_ecotype+shape:shore+size\_ratio:ref\_ecotype+shore:size\_ratio3}$ |
| 4251.8 | $\text{mountYN}\text{ \textasciitilde size\_ratio2+ref\_ecotype+log\_female:shore+shape:ref\_ecotype+size\_ratio:ref\_ecotype+shore:size\_ratio3}$ |
| 4252.4 | $\text{mountYN}\text{ \textasciitilde size\_ratio2+log\_female:ref\_ecotype+log\_female:shore+shape:ref\_ecotype+size\_ratio:ref\_ecotype+shore:size\_ratio3}$ |
| 4252.6 | $\text{mountYN}\text{ \textasciitilde size\_ratio+size\_ratio2+log\_female:shore+shape:ref\_ecotype+shape:shore+shore:size\_ratio3}$ |
| 4252.6 | $\text{mountYN}\text{ \textasciitilde size\_ratio2+log\_female:shore+shape:ref\_ecotype+size\_ratio:ref\_ecotype+ref\_ecotype:size\_ratio3+shore:size\_ratio3}$ |
| 4252.8 | $\text{mountYN}\text{ \textasciitilde log\_female:shore+shape:ref\_ecotype+shape:shore+size\_ratio:ref\_ecotype+size\_ratio2:ref\_ecotype+shore:size\_ratio3}$ |
| 4252.9 | $\text{mountYN}\text{ \textasciitilde ref\_ecotype+log\_female:shore+shape:ref\_ecotype+size\_ratio:ref\_ecotype+size\_ratio2:ref\_ecotype+shore:size\_ratio3}$ |

**Table S2.** Summary of parameter estimates for the top (lowest $\text{AIC}$) generalised linear models among $35442$ models. Maximum Likelihood Estimate and standard error ($\text{SE}$) of the variables and two-way interactions after full model average.

| Coefficients | Estimate | SE |
| --- | --- | --- |
| $\text{intercept}$ | $0.30$ | $0.35$ |
| $\text{size ratio}$ | $0.28$ | $0.67$ |
| $\text{size ratio}^{\boldsymbol{2}}$ | ${-3.00}^{**}$ | $0.94$ |
| $\text{size ratio}^{\boldsymbol{3}}$ | $0.12$ | $0.38$ |
| $\ln\left( \text{female size} \right)$ | $-0.05$ | $0.17$ |
| $\text{shape}$ | $-0.05$ | $0.37$ |
| $\text{reference wave}$ | $0.02$ | $0.08$ |
| $\text{size ratio : reference crab}$ | ${1.70}^{*}$ | $0.73$ |
| $\text{size ratio : reference wave}$ | $1.28$ | $0.68$ |
| $\text{size ratio}^{\boldsymbol{2}}\text{ : reference crab}$ | $-0.28$ | $0.91$ |
| $\text{size ratio}^{\boldsymbol{2}}\boldsymbol{:}\text{reference wave}$ | $-0.28$ | $0.92$ |
| $\text{size ratio}^{\boldsymbol{2}}\boldsymbol{:}\text{shape}$ | $-0.08$ | $0.43$ |
| $\text{size ratio}^{\boldsymbol{3}}\boldsymbol{:}\text{CZA}$ | ${1.09}^{*}$ | $0.51$ |
| $\text{size ratio}^{\boldsymbol{3}}\boldsymbol{:}\text{CZB}$ | $0.33$ | $0.51$ |
| $\text{size ratio}^{\boldsymbol{3}}\boldsymbol{:}\text{CZC}$ | $0.22$ | $0.50$ |
| $\text{size ratio}^{\boldsymbol{3}}\boldsymbol{:}\text{CZD}$ | $0.75$ | $0.51$ |
| $\text{size ratio}^{\boldsymbol{3}}\boldsymbol{:}\text{reference crab}$ | $0.06$ | $0.26$ |
| $\text{size ratio}^{\boldsymbol{3}}\boldsymbol{:}\text{reference wave}$ | $0.07$ | $0.30$ |
| $\ln\left( \text{female size} \right)\mathbf{:CZA}$ | $-0.41$ | $0.23$ |
| $\ln\left( \text{female size} \right)\boldsymbol{:}\mathbf{CZB}$ | $0.17$ | $0.23$ |
| $\ln\left( \text{female size} \right)\boldsymbol{:}\mathbf{CZC}$ | $0.30$ | $0.23$ |
| $\ln\left( \text{female size} \right)\boldsymbol{:}\mathbf{CZD}$ | $-0.40$ | $0.24$ |
| $\ln\left( \text{female size} \right)\text{: reference crab}$ | $-0.03$ | $0.11$ |
| $\ln\left( \text{female size} \right)\text{: reference wave}$ | $-0.02$ | $0.10$ |
| $\text{shape : reference crab}$ | $0.12$ | $1.51$ |
| $\text{shape : reference wave}$ | ${-3.29}^{*}$ | $1.62$ |
| $\text{shape : CZB}$ | $-0.81$ | $1.98$ |
| $\text{shape : CZC}$ | $-0.74$ | $1.83$ |
| $\text{shape : CZD}$ | $-1.14$ | $2.61$ |

Estimates followed by * indicate P-values $< 0.05$, ** indicate P-values $< 0.01$.

**HIERARCHICAL MODELS**

Our data allowed us to examine whether there were additional components to the mating pattern that were not captured by the size ratio, by adding to Eq. 1 information about covariates: sampling location (island), ecotype of the reference snail, shape and sex of the transect snails. The first covariate was included to test whether the mating pattern was conserved across experiments where variation could be due to the sampling location, the sampling time, or uncontrolled variation in experimental conditions between experiments. The second and the third ones were added to investigate any difference in mating pattern between ecotypes. The fourth covariate was used to check that the pattern was not influenced by the sex of the transect snail. This was achieved by replacing parameters in Eq. 1 by hyperparameters that are functions of the covariates, using hierarchical linear regression models in Stan.

The hierarchical configuration was adopted in a progressive way starting with one hyperparameter and one covariate and finishing with four hyperparameters and 16 covariates (four covariates per hyperparameter). For example, to test whether part of the variation in the mating pattern could be explained by different mating rates across the four islands, we built the $b_{1}$-island-hierarchical model where one parameter of Eq. 1, $b_{1}$, was defined as a ‘hyperparameter’ that was a function of one covariate, island, as follows:

$$b_{1}=\beta_{0}+\beta_{t}$$

(S1)

Here, $\beta_{0}$ is the intercept as well as the coefficient of the island CZA while $\beta_{t}$ is the effect, $t=1,2,3$ of the $\text{island}$ = CZB, CZC, CZD.

To examine instead whether the mating rate differed between ecotypes, we fitted the $b_{1}$-ecotype-hierarchical model which had the same hyperparameter as the model above but a different set of covariates: the ecotype of the reference snail and the shape of the transect snails. It was defined as:

$$b_{1}=\beta_{0}+\beta_{4}+\beta_{5}\text{T}$$

(S2)

Here, $\beta_{0}$ is the intercept as well as the coefficient of the reference ecotype Crab, $\beta_{4}$ is the effect of the reference ecotype Wave and $\beta_{5}$ is the regression coefficient associated with the shape of the transect snail (T).

The same logic was applied when testing for variation in the mating rate that could be explained by sex differences of the transect snails. The model $b_{1}$-sex-hierarchical contained one hyperparameter and one covariate and it was expressed as:

$$b_{1}=\beta_{0}+\beta_{6}.$$

(S3)

Here, $\beta_{0}$ is the intercept as well as the coefficient of the transect female snails and $\beta_{6}$ is the effect of the transect males.

Naturally, it was possible to add two or more covariates and test whether a combination of variables and their interactions (e.g., island and ecotype, island and sex) could explain any of the variation in the mating pattern in addition to female-male size ratios. These models were:

| $b_{1}\text{-island-ecotype-hierarchical}$ | $b_{1}=\beta_{0}+\beta_{t}+\beta_{4}+\beta_{5}\text{T}$. | (S4) |
| --- | --- | --- |
| $b_{1}\text{-island-sex-hierarchical}$ | $b_{1}=\beta_{0}+\beta_{t}+\beta_{6} .$ |  |
| $b_{1}\text{-ecotype-sex-hierarchical}$ | $b_{1}=\beta_{0}+\beta_{4}+\beta_{5}\text{T}+\beta_{6}+\beta_{6}\text{ : }\beta_{5}\text{T}$. |  |
| $b_{1}\text{-island-ecotype-sex-hierarchical}$ | $b_{1}=\beta_{0}+\beta_{t}+\beta_{4}+\beta_{5}\text{T}+\beta_{6}\text{e}+\beta_{6}\text{ : }\beta_{5}\text{T}$. |  |

Here, the colon ( : ) stands for the interaction between two variables.

As the hierarchical structure was developed from Eq. 1 (a non-hierarchical formulation), we took advantage of the inferred posterior distributions to define informative priors for the parameters of the hierarchical models. The prior of the intercept was set to the posterior distribution of the parameter $b_{1}$ of the non-hierarchical model and its inference was set to start from the $b_{1}$ mean posterior value. The priors of the regression coefficients were normally distributed with mean zero and standard deviation one, with starting value at zero. Finally, the priors of parameter $b_{0}$, $c$, $d$ and $\alpha$ were approximated to the posteriors of the non-hierarchical model and with starting values equivalent to the means of the non-hierarchical posterior distribution.

The same approach was applied to test hierarchical models for the other parameters, and to generate models with more than one hyperparameter. Hierarchical versions were not tested for parameter, $b_{0}$, which was fixed according to the mean of the posterior distribution estimated in the non-hierarchical model.

For each model, we computed the predictive accuracy using adjusted leave-one-out cross-validation (‘loo’ package in R, Vehtari et al. 2018) that returned the expected log predictive density (elpd, equivalent to the log-likelihood) and its standard error. Comparing pairs of models, the model with the higher elpd is the one that fits the data better (described in Vehtari et al. 2015, 2017). The elpd difference between two models was considered significant if it was larger than twice the estimated standard error.

**MODEL COMPARISONS**

The full hierarchical model was a better fit to the data than the non-hierarchical model, but not strongly so (elpd difference $=47.9$, $\text{SE}=10.7$; Table S3). This indicated that the mating pattern was influenced by factors other than the size ratio, although other effects were small.

**Table S3.** Comparison between the non-hierarchical and full hierarchical models. The estimate and standard error ($\text{SE}$) are reported for the expected log-predictive density (elpd loo), expected number of parameters (p loo) and the information criterion (looic $= -2*$ elpd loo).

| Non-hierarchical | Estimate | SE |  | Full hierarchical | Estimate | SE |
| --- | --- | --- | --- | --- | --- | --- |
| elpd loo | -2272.1 | 28.9 |  | elpd loo | -2224.2 | 27.1 |
| p loo | 4.9 | 0.5 |  | p loo | 4.7 | 0.1 |
| looic | 4544.2 | 57.9 |  | looic | 4448.5 | 54.2 |

To find out which parameters of the non-hierarchical model were most influenced by other explanatory variables, we investigated the posterior distributions of the parameters in the full hierarchical model. The strongest effects were for parameter $b_{1}$, mating rate, with some effects on the center, $c$. There was no effect on the ratio dependence, $d$, nor on the skew, $\alpha$, and so these two parameters were considered not to be influenced by other explanatory variables. The effects can be seen in the posterior distributions from the full model (Fig. S2).

We then considered which explanatory variables had the greatest influence on mating rate and center of the mating probability function. Again, we investigated the posterior distributions of the regression coefficients of the full hierarchical model and filtered out those explanatory variables associated with a regression coefficient whose posterior distribution was overlapping zero (Fig. S2). We were left with three different variables: island and sex of the transect snail for parameter $b_{1}$, mating rate, and island and ecotype of the reference snail for parameter $c$, the center. Another hierarchical model was built using only these variables with large effects to test whether it fitted the data as accurately as the full hierarchical model.

The hierarchical model based on the subset of variables with large effects fitted the data with no significant difference from the full hierarchical model (elpd difference $= 0.1$, $\text{SE}=5$; Table S4). This confirmed that the parameters of the non-hierarchical model that were most influenced by explanatory variables other than size ratio were indeed the mating rate, $b_{1}$, and the center, $c$. It also corroborated the inference that island and sex of the transect snail had the greatest effects on the mating rate and that island and the ecotype of the reference snail were the main explanatory variables for variation in the center of the mating function.

The mating rate was higher for the islands CZB and CZC than CZA and CZD and it was also higher for mating pairs where the transect snail was male (Fig. S3 left panel). The center of the skew normal mating distribution, and thus the OR, was shifted towards smaller values for the islands CZB and CZC than CZA and CZD. Similar shifts occurred when the reference snail belonged to the Wave ecotype. This means that OR was marginally closer to zero (female size = male size) in CZB and CZC compared to CZA and CZD and also in mating pairs with the Wave reference ecotype rather than Crab (Fig. S3 right panel).

Thus the mating pattern of *L. saxatilis* under the conditions of the no-choice experiment varied primarily in respect to the relative size between females and males of the mating pairs with marginal deviations in the mating rate due to other factors, primarily to island and sex of the transect snail. The center, hence OR, also varied marginally among islands (representing both sources of snails and time of the experiments) and ecotypes of the reference snail. The biological relevance of these minor variations was very weak because they mainly corresponded to a difference in mating probability rather than a distinction in the mating pattern. For this reason and for the technical and computational advantage of working with a simple model, we chose to perform the downstream analyses using the non-hierarchical model as it was the most meaningful model for quantifying the reduction in gene flow between Crab and Wave populations owing to the snail mating pattern.

**Table S4.** Comparison between the hierarchical model with the subset of variables and the full hierarchical model. The estimate and standard error ($\text{SE}$) are reported for the expected log-predictive density (elpd loo), expected number of parameters (p loo) and the information criterion (looic $= -2*$ elpd loo).

| Sub hierarchical | Estimate | SE |  | Full hierarchical | Estimate | SE |
| --- | --- | --- | --- | --- | --- | --- |
| elpd loo | -2224.4 | 29.1 |  | elpd loo | -2224.2 | 27.1 |
| p loo | 3.9 | 0.1 |  | p loo | 4.7 | 0.1 |
| looic | 4448.7 | 58.1 |  | looic | 4448.5 | 54.2 |

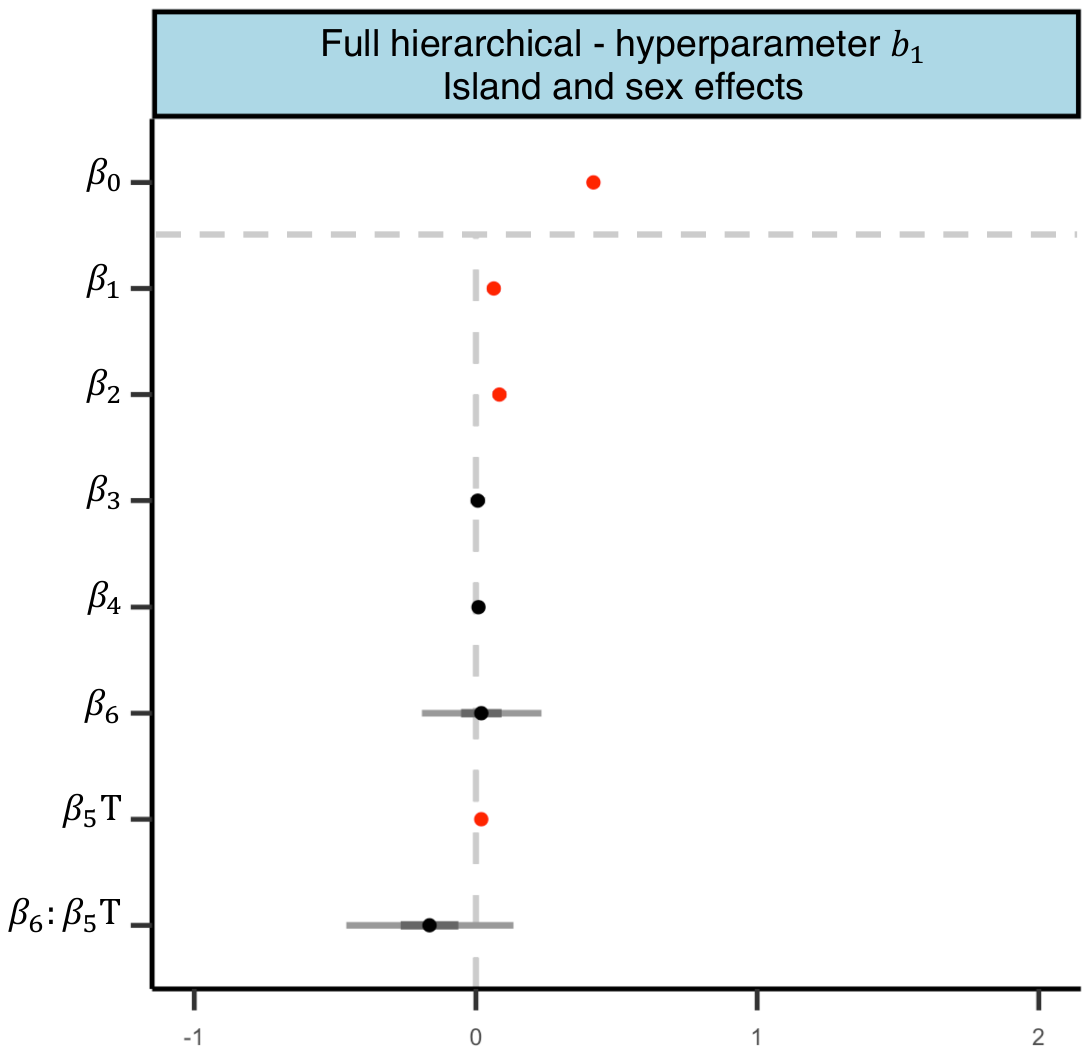

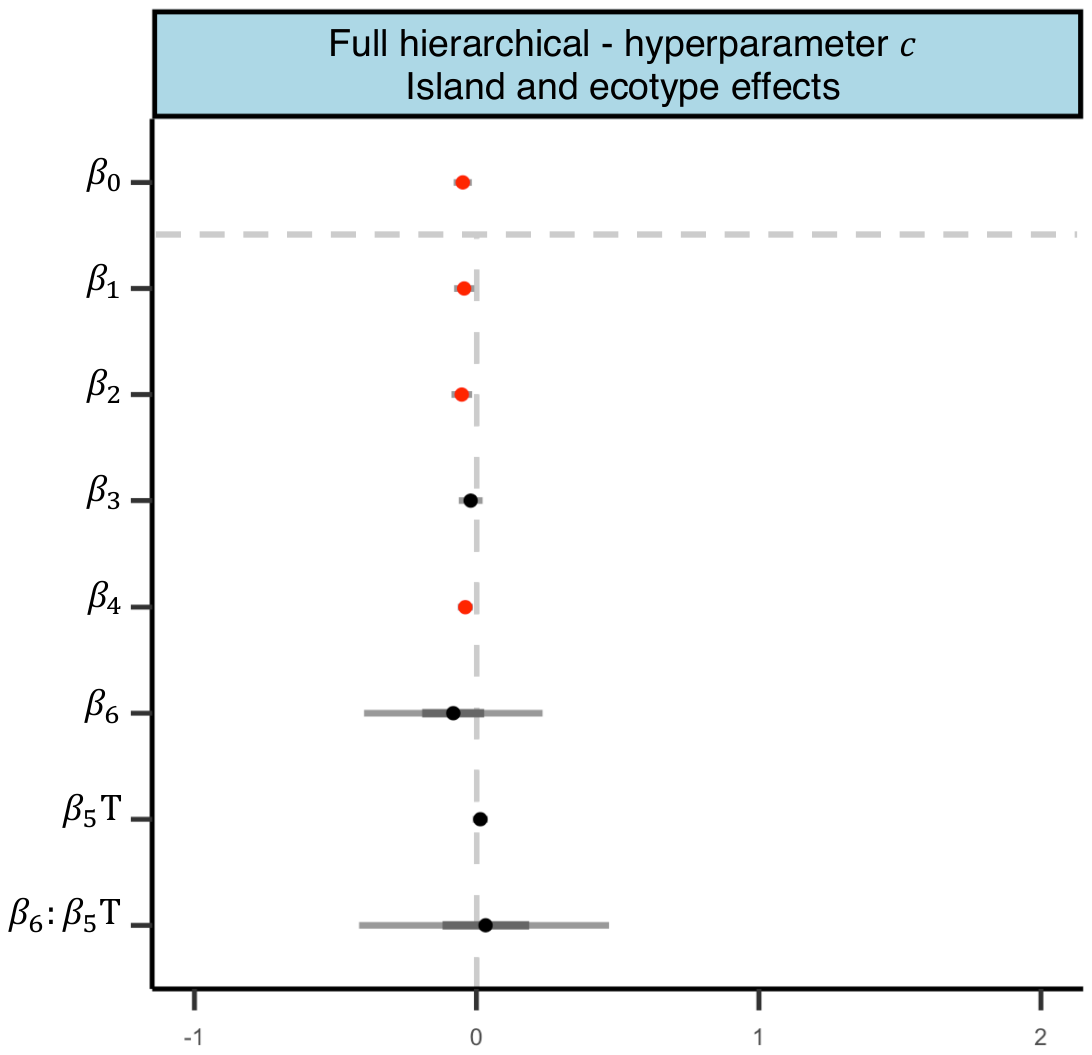

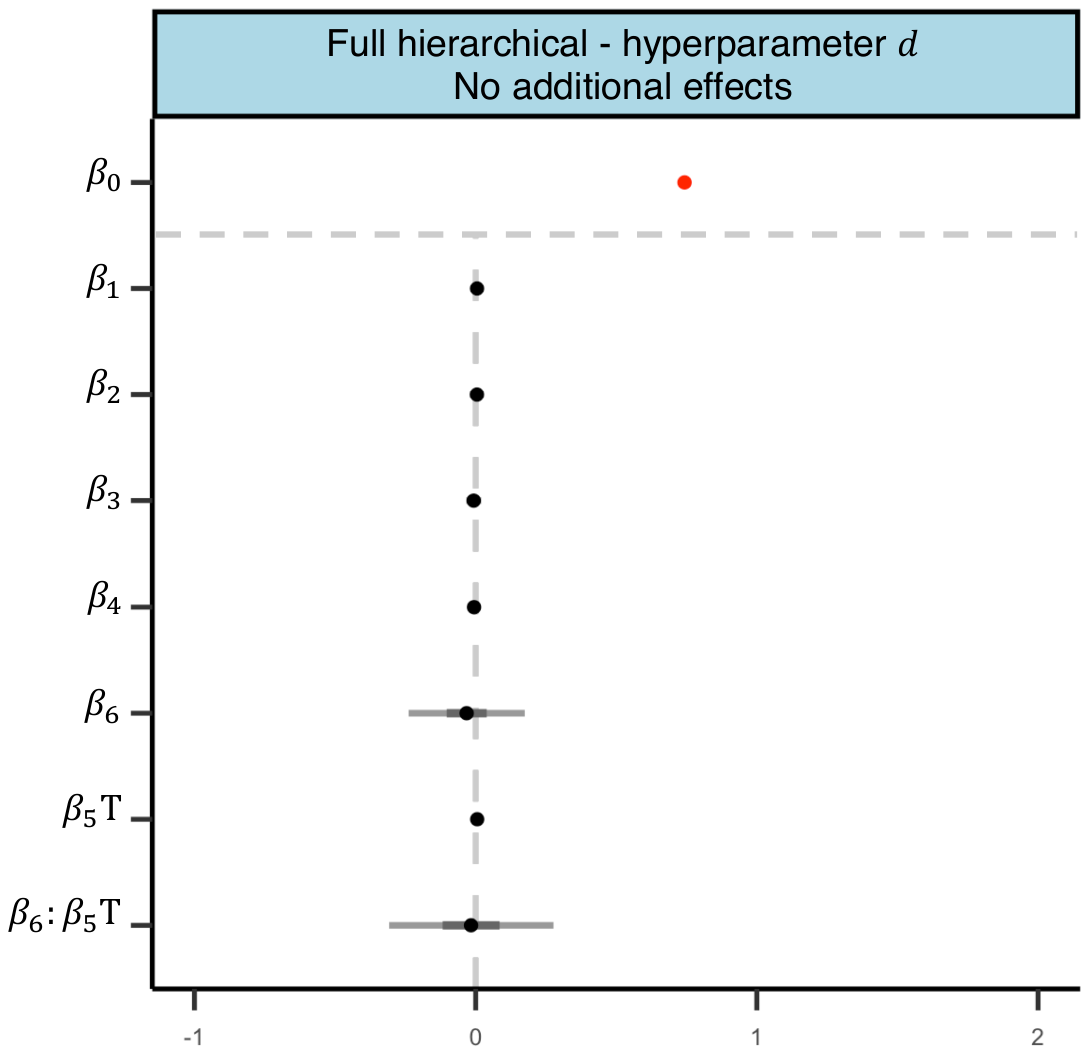

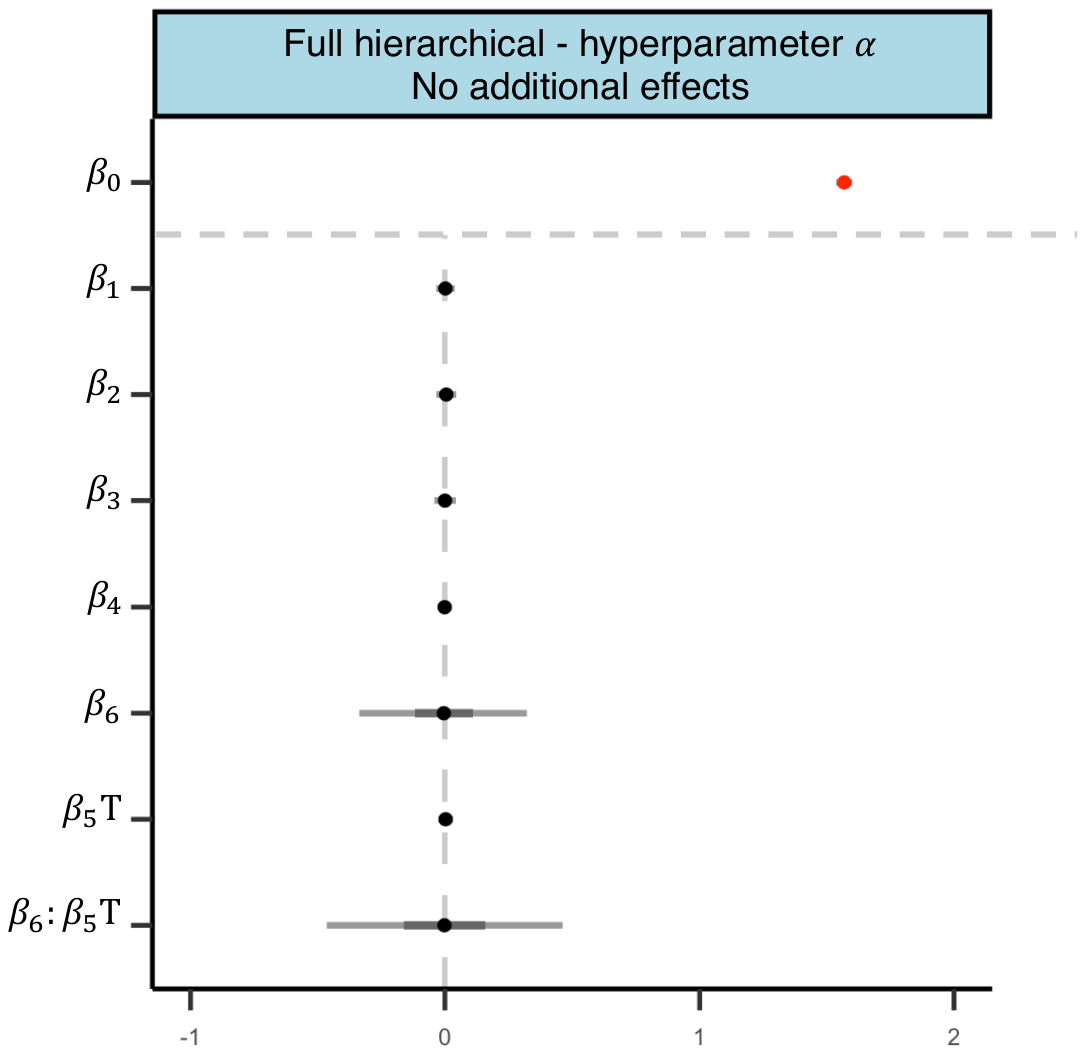

**Figure S2.** Posterior distributions of parameters in the full hierarchical model with four hyperparameters. Parameters (listed on the y axis) with mean (filled circle), 50% credible intervals (dark grey lines) and 95% credible intervals (light grey lines). The title of each panel contains the name of the hyperparameter and the covariate(s) with large effects. The posterior distribution of the intercepts, $\beta_{0}$, reflects the posterior distribution of the corresponding parameter estimated in the non-hierarchical model and any deviations from these parameters are illustrated by the posterior distributions associated with the covariates (island, ecotype and sex). The result is for one model which contains a total of 32 parameters (8 parameters for each of the four hyperparameters). Red circles indicate cases where the credible intervals do not include zero.

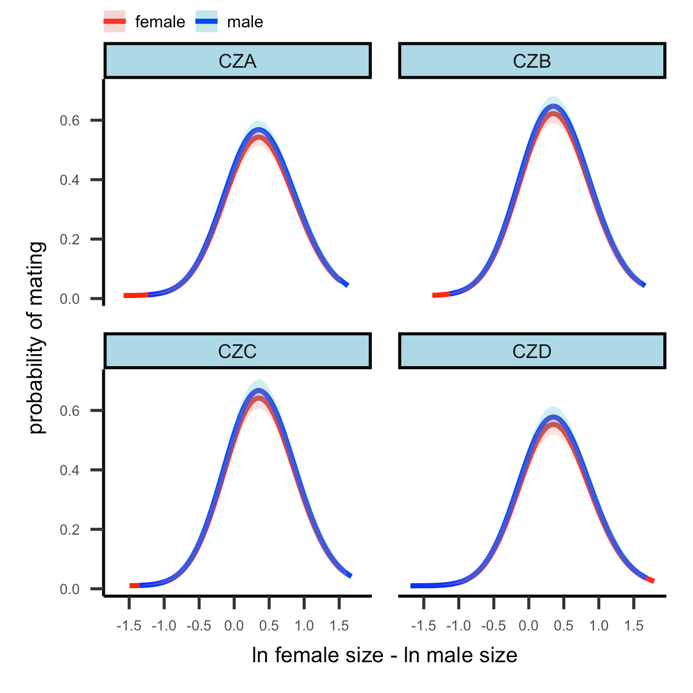

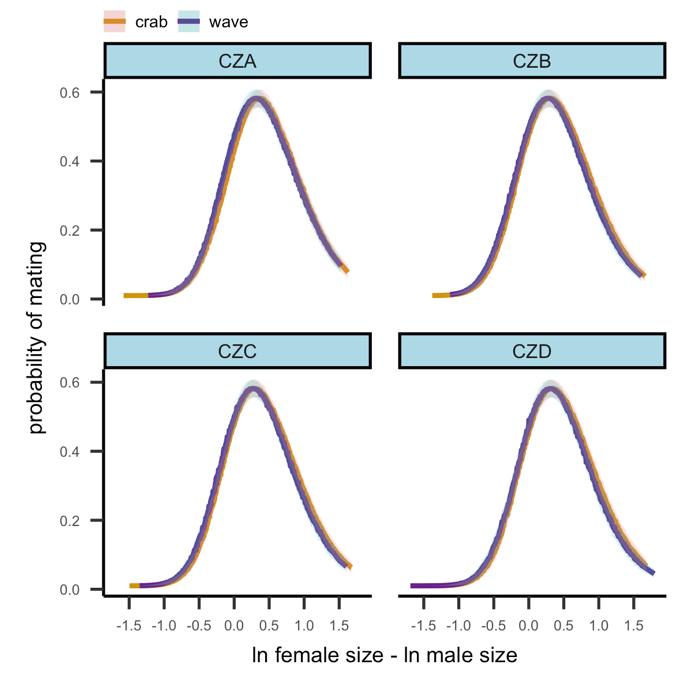

**Figure S3.** Hierarchical model fits for the variables with large effects on the mating rate and center. The probability of mating (y axis) was extracted from the hierarchical models with the highest expected log predictive density (left panel: $b_{1}$-island-sex-hierarchical model; right panel: $c$-island-ecotype-hierarchical model) and it was plotted against the size ratio (x axis) according to which covariates were included in the model (island effect: panel facets, sex of the transect snail: female in red and male in blue, ecotype of the reference: Crab in dark gold and Wave in dark purple).

**MATING PATTERN CONSEQUENCES IN THE CONTACT ZONE**

**Table S5.** Fitted cline parameter values from the hybrid zone analysis for ln(shell size). For each island and each side of the bays (left., right), maximum likelihood values [95% confidence intervals] estimated in the cline analysis are listed for the parameters on their respective scales. The standard deviations of ln(size) (SD) are reported for the crab and wave habitat as well as the SD at each of the two cline centers.

| **Parameter** | **CZA** | **CZB** | **CZC** | **CZD** |
| --- | --- | --- | --- | --- |
| $\text{left center} \text{(m)}$ | 140.1  [137.84-142.36] | 86.12 [85, 87.25] | 58.25 [58.06, 58.44] | 74.04 [72.2, 75.88] |
| $\text{right center} \text{(m)}$ | 279.74  [277.41, 282.06] | 143.57  [139.72, 147.43] | 128.75  [126.73, 130.77] | 181.8  [179.97, 183.64] |
| $\text{left width (m)}$ | 31.69  [26.74, 37.56] | 5.39 [3.77, 7.71] | 3.43 [6.85, 1.72] | 15.64  [12.18, 20.09] |
| $\text{right width (m)}$ | 22.94 [18.08, 29.11] | 21.85 [13.63, 35.02] | 19.18 [15.49, 23.76] | 9.42 [6.24, 14.22] |
| $\text{crab male (mm)}$ | 11.28 [11.46, 11.1] | 11.28 [11.6, 10.97] | 10.83  [11.07, 10.59] | 11.16  [11.38, 10.94] |
| $\text{wave male (mm)}$ | 4.4 [4.61, 4.19] | 5.25 [5.49, 5.03] | 3.77 [3.93, 3.61] | 4.22 [4.49, 3.98] |
| $\text{crab female (mm)}$ | 11.94 [12.18, 11.7] | 11.59 [12.06, 11.13] | 11.25 [11.7, 10.8] | 11.7 [12.18, 11.25] |
| $\text{wave female (mm)}$ | 5.16 [5.58, 4.76] | 5.93 [6.3, 5.58] | 4.81 [5.1, 4.53] | 5.26 [5.81, 4.76] |
| $\text{crab SD} \text{(mm)}$ | 1.13 [1.11, 1.14] | 1.11 [1.09, 1.12] | 1.16 [1.14, 1.17] | 1.12 [1.1, 1.13] |
| $\text{left center SD} \text{(mm)}$ | 1.31 [1.24, 1.39] | 1.52 [1.37, 1.7] | 1.21 [1.08, 1.58] | 1.42 [1.3, 1.55] |
| $\text{right center SD} \text{(mm)}$ | 1.46 [1.35, 1.58] | 1.45 [1.32, 1.59] | 1.58 [1.47, 1.71] | 1.77 [1.5, 2.08] |
| $\text{wave SD} \text{(mm)}$ | 1.4 [1.37, 1.43] | 1.39 [1.37, 1.42] | 1.34 [1.31, 1.37] | 1.47 [1.43, 1.51] |

**ASSORTATIVE MATING AND SEXUAL SELECTION**

**Figure S4.** Assortative mating and sexual selection in the CZA, CZC and CZD transects. Habitat boundaries are marked by black vertical dashed lines, the crab habitat is the region inside (grey fill) and the wave habitat is outside (white fill) the two dashed lines. Cline facet: $\ln\left( \text{size} \right)$ of transect snails in bins (dots with $95\% \text{CI}$) and fitted clines (solid lines $\pm\text{SD}$) for females (in red) and males (in blue). AM facet: strength of assortative mating measured as the Pearson correlation coefficient ($r$) between female and male $\ln\left( \text{size} \right)$ of mated pairs. DSS facet: directional component of sexual selection measured as the difference in mean $\ln\left( \text{size} \right)$ of mated males compared to mated plus non-mated males. The black horizontal dashed line indicates where this component is absent. SSS facet: stabilizing component of sexual selection calculated as the difference in variance between mated male $\ln\left( \text{size} \right)$ and mated plus non-mated male $\ln\left( \text{size} \right)$. The black horizontal dashed line indicates where this component is absent.

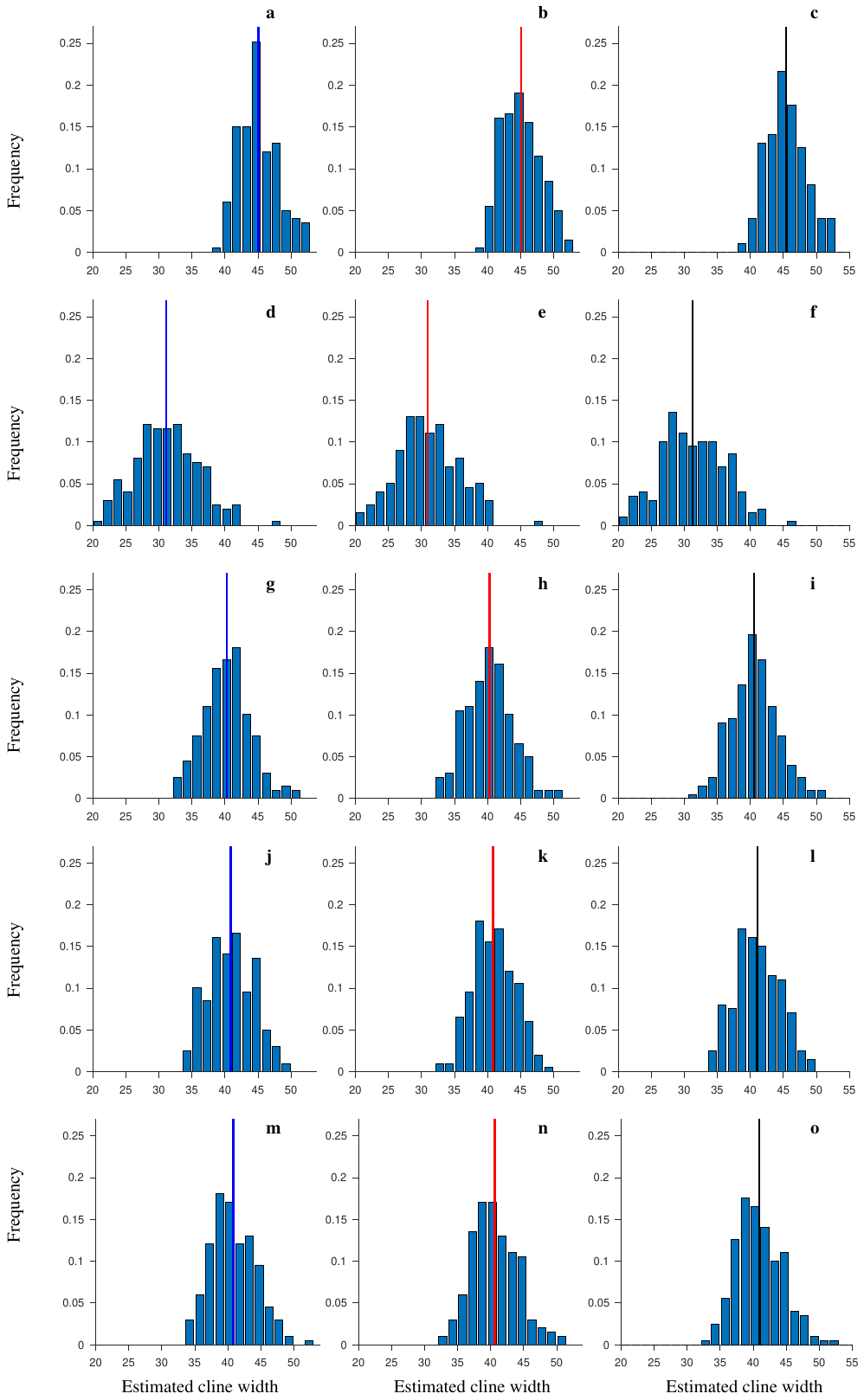

**Figure S5.** The distribution of estimated cline widths for the hybrid index, at the end of the simulations, for five models of mating: *RM* (a-c), *AS* d-f), *SimMR* (g-i), *SimOR* (j-l), And *SimM0* (m-o). We show the results for males only (first column), females only (second column), and all individuals together (third column). Vertical lines show the mean values. $200$ independent realizations of each model.

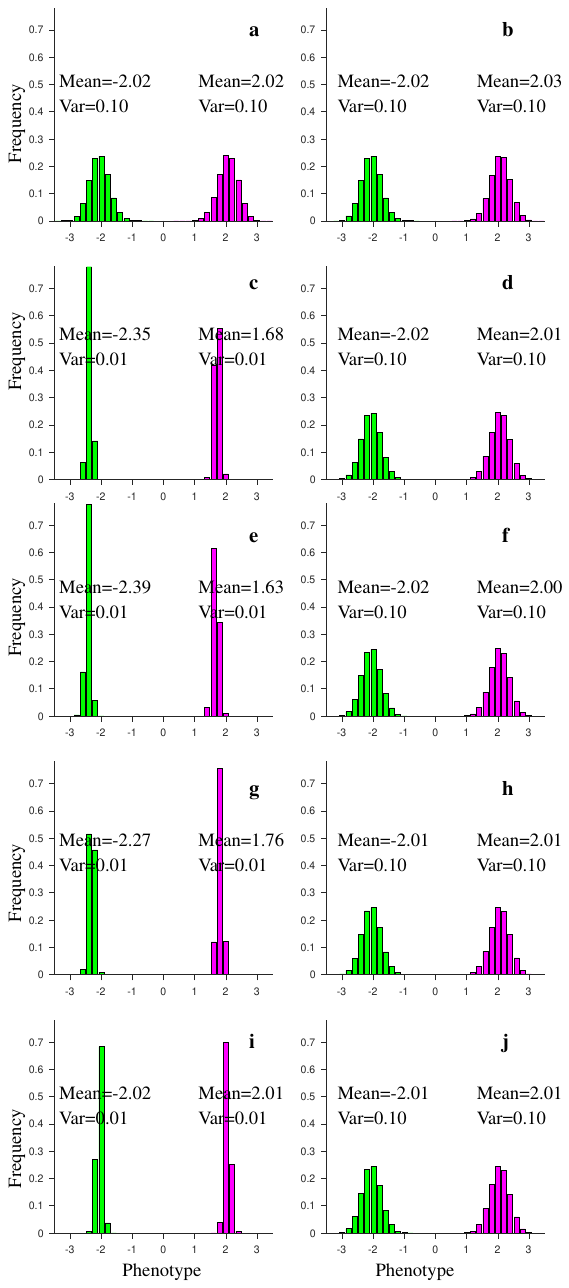

**Figure S6.** Distribution of phenotypes at the two habitat ends at the end of the simulations for *RM* model (a-b), *AS* model (c-d), *SimMR* model (e-f), *SimOR* model (g-h), and *SimM0* model (i-j). The results for males are shown in the first column (a, c, e, g, i), and for females in the second column (b, d, f, h, j). In each panel, we show the distribution of phenotypes from the 30 right-most patches in the habitat (green, distributions on the left), and from the 30 left-most patches in the habitat (magenta, distributions on the right). The observed variation is a balance between gene flow, segregation and stabilizing selection in each case. Narrower distributions reflect stronger selection while the means reflect the compromise between natural and sexual selection. Above each distribution we give mean and variance. Distributions are averaged over 200 independent realizations of each model.
